# Next generation vaccine platform: polymersomes as stable nanocarriers for a highly immunogenic and durable SARS-CoV-2 spike protein subunit vaccine

**DOI:** 10.1101/2021.01.24.427729

**Authors:** Jian Hang Lam, Amit Kumar Khan, Thomas Andrew Cornell, Regine Josefine Dress, Teck Wan Chia, Wen Wang William Yeow, Nur Khairiah Mohd-Ismail, Shrinivas Venkatraman, Kim Tien Ng, Yee-Joo Tan, Danielle E. Anderson, Florent Ginhoux, Madhavan Nallani

## Abstract

Multiple successful vaccines against SARS-CoV-2 are urgently needed to address the ongoing Covid-19 pandemic. In the present work, we describe a subunit vaccine based on the SARS-CoV-2 spike protein co-administered with CpG adjuvant. To enhance the immunogenicity of our formulation, both antigen and adjuvant were encapsulated with our proprietary artificial cell membrane (ACM) polymersome technology. Structurally, ACM polymersomes are self-assembling nanoscale vesicles made up of an amphiphilic block copolymer comprising of polybutadiene-b-polyethylene glycol and a cationic lipid 1,2-dioleoyl-3-trimethylammonium-propane. Functionally, ACM polymersomes serve as delivery vehicles that are efficiently taken up by dendritic cells, which are key initiators of the adaptive immune response. Two doses of our formulation elicit robust neutralizing titers in C57BL/6 mice that persist at least 40 days. Furthermore, we confirm the presence of memory CD4^+^ and CD8^+^ T cells that produce Th1 cytokines. This study is an important step towards the development of an efficacious vaccine in humans.

## Introduction

Vaccines are an integral part of the global healthcare strategy and have played a decisive role in eliminating or controlling numerous infectious diseases. Since coronavirus disease 2019 (Covid-19) emerged as a pandemic, rapid development of a vaccine has become a paramount focus across the globe. The etiological agent, severe acute respiratory syndrome coronavirus 2 (SARS-CoV-2), is capable of efficient human-to-human transmission (1-3), with ∼20% of patients exhibiting severe (respiratory distress) to critical (respiratory failure, septic shock and/or multi organ failure) symptoms (4). SARS-CoV-2 belongs to the genus *Betacoronavirus* within the family *Coronaviridae* (5). Each virion consists of a nucleocapsid protein-encapsulated single-stranded genomic RNA, surrounded by a lipid bilayer into which spike (S), membrane and envelope proteins are incorporated (6). Trimers of S protein form spike-like projections from the virus exterior surface and are key to host-virus interaction. The S protein, which consists of subunits S1 and S2, enables viral entry into the host cell through the interaction of the receptor binding domain (RBD; situated within the S1 subunit) with the angiotensin-converting enzyme 2 (ACE2) receptor of the host cell membrane. This stimulates cleavage at the S1-S2 junction by host cell proteases and induces significant structural rearrangement that exposes the hydrophobic fusion peptide, thus permitting the merging of viral and host cell membranes leading to viral entry (7). The spike protein is immunogenic and the target of antibodies as well as T cells, particularly CD4^+^ T cells (8-10). Therefore, it has emerged as the key target for subunit vaccines of various modalities.

Meeting the global demand for a Covid-19 vaccine using traditional approaches of inactivated or live attenuated virus is challenging due to the requirement for biosafety level (BSL) 3 facility to handle SARS-CoV-2. Subunit vaccines based on the spike protein eliminate the need for handling live virus and are key to addressing the global demand challenge. Advances in structural biology, development of specialized carriers for respective cargoes (including mRNA and protein), coupled with the rapid dissemination of the SARS-CoV-2 genomic sequence, have greatly accelerated the development of subunit vaccines.

Several candidates are in clinical trials and a few are seeking regulatory approval for emergency use (11). Although the clinical data has been promising, some of the leading vaccine candidates do possess significant limitations. Adenoviral vectors could trigger anti-vector responses that may reduce the efficacy of subsequent administrations (12). mRNA vaccines formulated in lipid nanoparticles have enabled swift response to the Covid-19 pandemic but issues of stability (currently mitigated by an ultra-cold chain to preserve mRNA integrity) and cost pose a major hurdle for effective and equitable distribution of such vaccines (13).

Advancement in nanotechnology can potentially contribute to the development of a safe, cost-effective and scalable vaccine platform, thus addressing some of the issues with the current Covid-19 vaccine candidates. Amphiphilic block copolymer self-assembly offers a straightforward, scalable route to well-defined nanoscale vesicles. By controlling the ratio of the different constituent blocks, self-assembly can be tailored to access different nanostructures, including polymersomes. The ability to compartmentalize antigens and adjuvants in the aqueous compartment of polymersomes renders them very attractive for vaccine application (14). Compared to liposomes, polymersomes have the unprecedented advantage of tuning membrane thickness and property (15). Owing to their relatively long hydrophobic segments (16, 17), polymersomes possess enhanced stability without the need for additional stabilization strategies such as cross-linking chemistries (18). In spite of their tremendous potential, only a few reports are available employing polymersomes as a carrier for vaccine application (19-21). Nevertheless, these limited studies clearly demonstrate that antigen-loaded polymersomes can target dendritic cells (DCs), the most efficient of antigen-presenting cells (APCs). Moreover, many polymersome attributes, such as size and surface properties, can be customized (17) to modulate their specific uptake by DCs, hence rendering polymersomes as an ideal platform for the delivery of subunit vaccines.

In the present work, we describe the development of a subunit vaccine based on the spike protein of SARS-CoV-2, co-administered with CpG adjuvant. We can encapsulate different classes of biomolecules (i.e.: DNA and protein) within our proprietary ACM polymersomes to produce coherent and immunogenic particles, thus demonstrating the flexibility of this technology. Structurally, ACM polymersomes are self-assembling vesicles made of an amphiphilic block copolymer comprising of polybutadiene-b-polyethylene glycol (PBD-PEO) and a cationic lipid 1,2-dioleoyl-3-trimethylammonium-propane (DOTAP) (22). We show here that functionally, ACM polymersomes serve as delivery vehicles which are efficiently taken up by DC1 and DC2, which are key initiators of the adaptive immune response. We further investigate the immunological effect of ACM polymersomes on different SARS-CoV-2 spike proteins, namely the ectodomain of the spike protein, the S2 domain only, and a trimeric spike protein. Altogether, we show that our vaccine formulation possesses strong immunogenicity and can elicit robust and durable humoral and cellular immunity against SARS-CoV-2 in C57BL/6 mice that persist for at least 40 days.

## Results

### Spike protein purification and encapsulation in ACM polymersomes

The SARS-CoV-2 spike protein is immunogenic and targeted by T cells and strongly neutralizing antibodies (8-10), making it a highly attractive subunit vaccine target. Based on previous work with various viral and cancer antigens (data not shown), we established that immunogenicity of a protein could be significantly improved through encapsulation within ACM polymersomes. Moreover, a further increase in the immune response could be achieved via co-administration of an appropriate adjuvant, such as the toll-like receptor (TLR) 9 agonist CpG. Therefore, our present approach involved the encapsulation of both the spike protein as well as CpG adjuvant for co-administration (Fig. 1a). To generate the spike protein, we engineered T.ni cells to express a spike variant that retained S1 and S2 domains but excluded the hydrophobic transmembrane domain (hereby referred to as “S1S2”; Fig. 1b), thereby improving protein solubility. In addition, we purchased a S2 fragment and a trimeric spike protein (Fig. 1b) from commercial vendors to serve as controls for the subsequent immunogenicity study. S2 was ideal as a negative control since it lacked strongly neutralizing epitopes whereas trimeric spike was used as a positive control given that it best represented the natural configuration of this viral protein.

**Figure 1.**
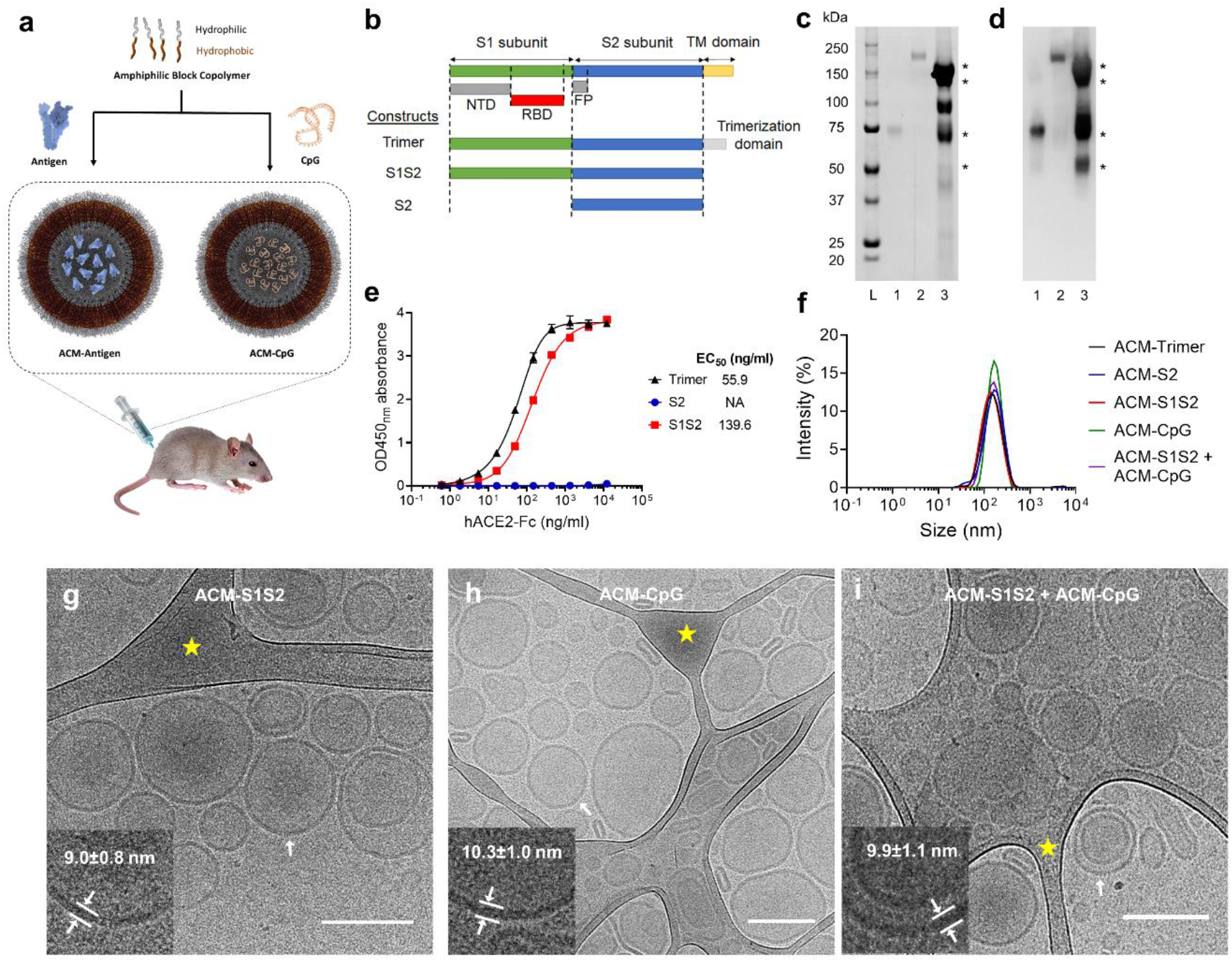
ACM-vaccine characterization. **a**. Schematic illustration of ACM-vaccine preparation. Antigens and CpG adjuvant were encapsulated within individual ACM polymersomes. A 50:50 v/v mixture of ACM-Antigen and ACM-CpG was administered to mice as the final vaccine formulation. **b**. Schematic of the spike protein variants used in this study. S1S2 protein was expressed and purified inhouse whereas S2 and trimer were purchased from commercial vendors. NTD: N-terminal domain. RBD: receptor binding domain. FP: fusion peptide. TM: transmembrane. **c**. SYPRO Ruby total protein stain. Lane L: Precision Plus Protein Standards (Bio-Rad). Lane 1: S2. Lane 2: trimer. Lane 3: S1S2. **d**. Western blot using mouse immune serum raised against SARS-CoV-2 spike. Western blot-reactive S1S2 bands are indicated by *. **e**. ACE2 binding curves of trimer, S2 and S1S2. **f**. Dynamic Light Scattering (DLS) measurements of ACM-antigens (ACM-trimer, ACM-S2 and ACM-S1S2), and ACM-CpG. ACM particles were determined to be 100-200 nm in diameter. **g-i**. Cryo-EM images of ACM-S1S2, ACM-CpG, and mixture of ACM-S1S2 + ACM-CpG illustrate the vesicular architecture with an average diameter of 158±25 nm (scale bar 200 nm). Inserts (lower left of each image) are magnifications of the bilayer membrane of vesicles at regions indicated by white arrows. Areas highlighted by yellow star are lacy carbon.

The three spike variants were analysed by SDS-PAGE followed by SYPRO Ruby staining (Fig. 1c) and western blot using mouse immune serum raised against a recombinant SARS-CoV-2 spike protein purchased from Sino Biological (Fig. 1d). Total protein staining using SYPRO dye showed S1S2 protein to consist of several bands, including two closely migrating major bands at the 150 kDa position, as well as two smaller bands at 75 kDa and 50 kDa (Fig. 1c). All four bands were recognized by spike-specific antibodies in western blot (Fig. 1d), confirming that they were all or parts of the spike protein. Among the two bands at the 150 kDa position, the heavier one corresponded to a highly glycosylated full-length spike protein, whereas the lighter one was presumed to have a lighter glycosylation profile. The remaining two western blot-reactive bands were likely truncations of the full-length protein. Interestingly, analytical size exclusion chromatography data indicated that our S1S2 protein could form higher order structures (311 kDa; Supplementary Fig. 1). This was larger than an expected monomer (180 kDa; (23)) and may suggest the presence of oligomers despite the absence of a trimerization domain. Functionally, our S1S2 protein bound ACE2 strongly with an EC50 value of 139.6 ng/ml (Fig. 1e) though its avidity was lower compared to trimeric spike.

Taken together, our data suggests a correctly folded spike protein that presents a functional receptor binding domain (RBD). Adopting the correct conformation is fundamentally important from an immunization standpoint since potently neutralizing antibodies typically target the RBD (9), though other regions of the spike protein have also been reported (8, 9, 24). In the scenario where vaccination is done using an incorrectly folded protein, induction of a high ratio of binding to neutralizing antibody is expected and may predispose the individual to antibody dependent enhancement (ADE) or vaccine-associated enhanced respiratory disease (VAERD) during actual infection (25). ADE describes a scenario in which binding but non-neutralizing antibodies facilitate viral entry into cells bearing Fcγ receptors, notably cells of the myeloid lineage, resulting in increased viral load and disease severity (26). With regards to VAERD, this refers to a situation wherein high levels of non-neutralizing antibodies causes excessive immune complex deposition, complement activation and, ultimately, airway inflammation (27, 28). Although both phenomena remain theoretical possibilities in the context of Covid-19, they still argue strongly for a conformationally-correct spike protein to enhance vaccine safety and efficacy.

ACM polymersomes are self-assembled from an amphiphilic block copolymer comprising of polybutadiene-b-polyethylene glycol (PBD-PEO) and a cationic lipid DOTAP (22). When amphiphilic block copolymers encounter water, they undergo self-assembly into a thermodynamically stable bilayer conformation placing the PBD block on the inside to minimise the water interaction, whereas the water-soluble PEG block is surface-exposed. During the self-assembly process, solutes are entrapped into the vesicular cavity. DOTAP, or other charged lipids, can be used to increase electrostatic interactions to enhance encapsulation of negatively charged molecules, such as antigens and adjuvants. In this work, viral antigens (spike trimer, S2 and S1S2 protein) and CpG adjuvant were separately encapsulated in individual vesicles as ACM-trimer, ACM-S2, ACM-S1S2 and ACM-CpG, respectively. Vesicles were extruded to within 100-200 nm diameter range followed by dialysis to remove the solvent, non-encapsulated antigens and adjuvant. The final vaccine formulation was a 50:50 v/v mixture of ACM-S1S2 and ACM-CpG prior to administration.

All samples were tested negative for endotoxin using colorimetric HEK Blue cell-based assay (Supplementary Fig. 2).

The sizes and morphologies of ACM-antigen and ACM-CpG were assessed by dynamic light scattering (DLS) and cryogenic-transmission electron microscopy (cryo-TEM), respectively. Overall, the sizes of ACM polymersomes were uniform (Fig. 1f) and followed a unimodal intensity-weighted distribution with a mean z-average hydrodynamic diameter of 158±25 nm. The sizes of the different ACM-antigen preparations were comparable – ACM-trimer: 133 nm (PDI 0.192); ACM-S1S2: 139 nm (PDI 0.181); and ACM-S2, 143 nm, (PDI 0.178). ACM-CpG, on the other hand, was slightly larger at 183 nm (PDI 0.085). The final vaccine formulation (ACM-S1S2 + ACM-CpG) showed a size distribution comparable with those of individual vesicles (Fig. 1f). Electron micrographs revealed a vesicular architecture with a homogeneous size distribution, suggesting topographically uniform vesicles (Fig. 1g-i). From line profile measurements, the bilayer thickness of ACM-S1S2, ACM-CpG, and ACM-S1S2 + ACM-CpG were estimated to be 9.0±0.8 nm, 10.3±1.0 nm and 9.9±1.1 nm, respectively.

To assess protein encapsulation within vesicles, ACM-antigen particles were lysed with 2.5% non-ionic surfactant Triton X100 and then characterized by SDS-PAGE alongside free protein calibration standards. The concentrations of encapsulated proteins were quantified by the densitometric method from SDS-PAGE followed by SYPRO Ruby staining (Supplementary Fig. 3a-c). ACM polymers interacted with SYPRO stain to generate a distinct smear at the bottom of the lane and co-localization of the protein band with this smear confirmed that encapsulation had occurred. The amounts of encapsulated trimer, S1S2 and S2 were determined to be 48 µg/ml, 46 µg/ml and 25.7 µg/ml, respectively, from 100 µg/ml starting concentrations. To remove free protein that escaped encapsulation, all ACM-preparations were dialyzed. A parallel dialysis experiment with free protein control was performed to determine the quantity of free protein remaining in each ACM preparation. SYPRO staining showed 19.8 µg/ml free trimer, 7.5 µg/ml free S1S2 protein and 0 µg/ml free S2 remaining after dialysis from 100 µg/ml starting protein concentrations (Supplementary Fig. 3a-c), indicating that majority of the non-encapsulated proteins were removed from ACM-S1S2 and ACM-S2 preparations, whereas close to 40% free protein still remained with the ACM-trimer sample. The lower efficiency of trimer removal may be caused by its larger size relative to S1S2 or S2, thus reducing its diffusion across the dialysis membrane. To quantify the concentration of CpG encapsulated in ACM vesicles, the DNA binding dye SYBR Safe was used. Based on the 530 nm fluorescent emission, the encapsulation of ACM-CpG was determined to be 480 µg/ml at an efficiency of 60%.

Given the importance of shelf life and product stability in the context of local and global distribution, we performed a stability study on free S1S2 protein, ACM-S1S2, free CpG, ACM-CpG, free S1S2 + free CpG and ACM-S1S2 + ACM-CpG at 4 °C and 37 °C. The initial observation showed a very stable vesicle with no change of size and PDI of the ACM-S1S2 formulation, no degradation of S1S2 protein content, and minimal loss of activity for up to 20 weeks at 4°C measured by DLS, SDS-PAGE followed by SYPRO staining, and ACE2 binding assay by ELISA, respectively (Supplementary Fig. 4a-d). However, an accelerated stability study at 37°C showed a decrease in protein concentration for both free S1S2 as well as ACM-S1S2 after one week (Supplementary Fig. 5a), indicating proteolytic degradation at elevated temperature. Unexpectedly, samples containing CpG (either ACM-S1S2 + ACM-CpG or free S1S2 + free CpG) exhibited reduced protein degradation. Further, only ACM-S1S2 + ACM-CpG maintained its protein content for up to 28 days, whereas other formulations showed complete proteolysis (Supplementary Fig. 5a). It remained unclear how CpG was able to maintain protein stability at 37°C, though we speculated that the negatively charged CpG may possibly associate with proteases present as impurities in the S1S2 sample, thereby hindering proteolysis of S1S2 protein. In contrast, the size and PDI of the ACM formulations remained stable over the 28-day time course (Supplementary Fig. 5b, c).

In summary, we had expressed and purified functional SARS-CoV-2 spike (“S1S2”) protein from T.ni cells that bound ACE2 with high avidity. This suggested a correctly folded protein, which was necessary for the induction of neutralizing antibodies. The protein and CpG adjuvant were separately encapsulated in ACM-polymersomes for the purpose of co-administration in our final vaccine formulation. In stability tests, the ACM-encapsulated S1S2 protein quickly degraded at 37°C but remained intact for at least 20 weeks at 4°C. With proper temperature control at 4°C during storage, transport and distribution, our ACM-S1S2 formulation would be expected to maintain functionality for prolonged periods.

### ACMs are efficiently taken up by dendritic cells

To gain long-lasting immunity against viruses, such as SARS-CoV-2, the initial induction of an efficient immune response is crucial. Dendritic cells (DCs), which can be roughly divided into XCR1/CD8/CD103^+^CD11b^-^ classical DC1 and CD11b^+^XCR1/CD8/CD103^-^ classical DC2 (29, 30), are cells of the innate immune system highly specialized in priming and activation of naïve CD8^+^ T cells and CD4^+^ T cells. Following the uptake of foreign antigens, DCs process and present them to naïve CD8^+^ T cells or CD4^+^ T cells via their MHC-I or MHC-II complexes, resulting in antigen-specific cytotoxic and helper T cell responses, respectively (30, 31).

To analyze the capacity of DCs to take up ACM polymersomes, mice were injected subcutaneously with either PBS, the non-toxic dye Rhodamine incorporated in ACMs or Rhodamine and DQ-OVA co-encapsulated in ACMs. One to six days post injection, expression of Rhodamine and/or DQ by DCs in the skin and skin-draining lymph nodes (LNs) was examined by flow cytometry. Both DC1 and DC2 could efficiently take up ACM-Rhodamine, as evidenced by the high Rhodamine signals recorded in both cell types (Fig. 2a, b). However, DC2 in the skin seemed to be more potent in ACM-Rhodamine uptake over time as compared to DC1, with around 80 - 90% of DC2 remaining Rhodamine^+^ on day three post injection compared to 80% and ∼20% of DC1 on days one and three, respectively (Fig. 2a, c). DC2 could also efficiently co-express Rhodamine and DQ-OVA, while DC1 seem to either have a lower take up rate or completely fail to take up ACMs co-encapsulated with Rhodamine and DQ-OVA (Fig. 2a, c). While we could not detect any signal for Rhodamine nor DQ-OVA in the draining LNs on day one post injection (data not shown), around two to five percent of migratory DC1 (mDC1) and migratory DC2 (mDC2) had taken up Rhodamine and/or Rhodamine and DQ-OVA, respectively, as early as day three post injection (Fig. 2b, d). These results show that DCs take up ACMs from the skin to then migrate to the draining LNs with their “ACM cargo”. As the LNs are the prime location for DC-T cell interaction, we speculate that these ACM-loaded DCs interact with naïve CD8^+^ T cells or CD4^+^ T cells to prime and polarize them for further responses.

**Figure 2.**
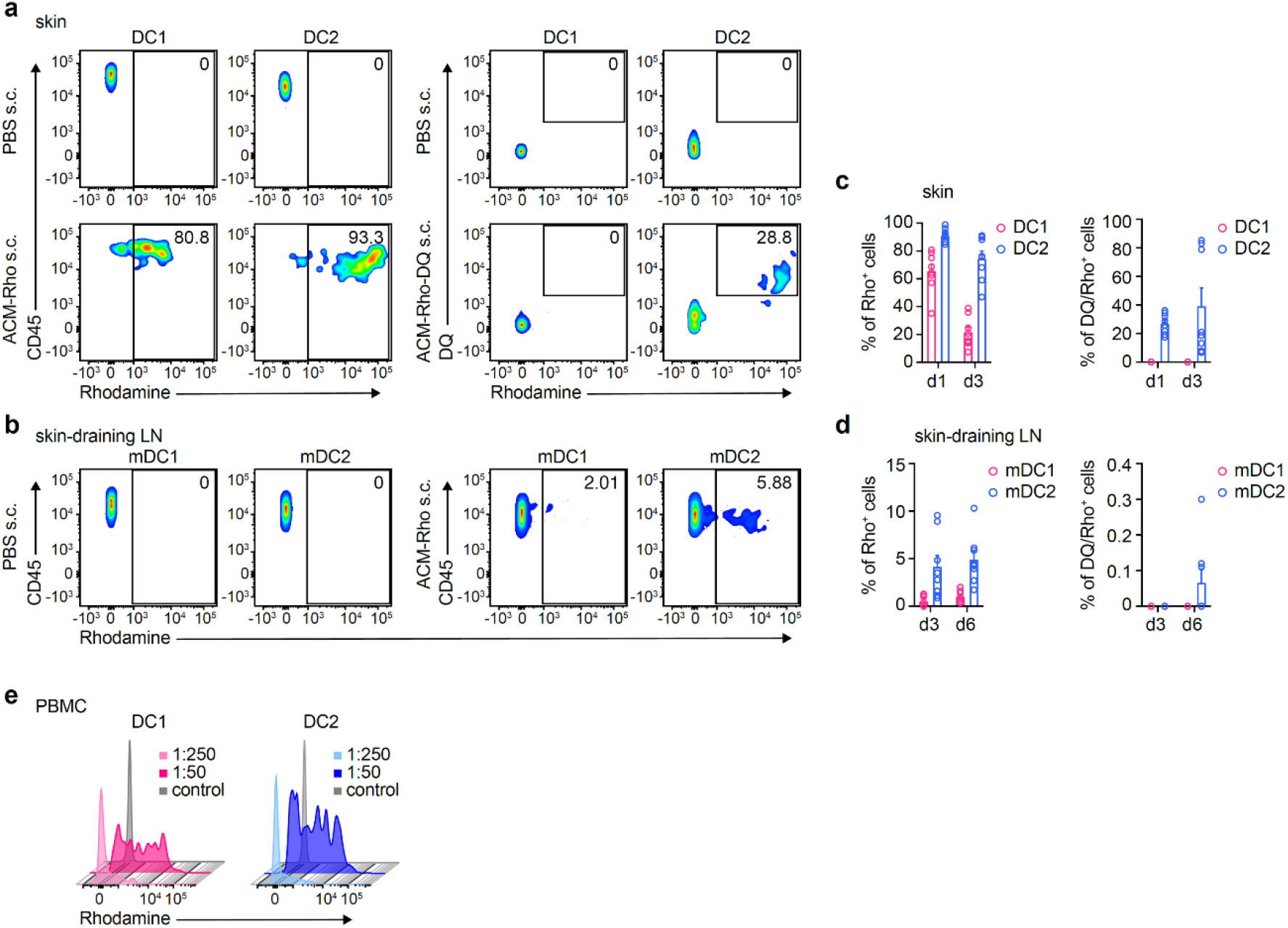
ACM vesicles were efficiently taken up by dendritic cells. **a-d**. Mice (n = 8) SC injected with ACM-Rho or ACM-DQ-OVA vesicles. **a**. Rho and Rho vs DQ signals from skin DC1 and DC2 one day post SC injection. **b**. Rho signals from migratory DC1 and DC2 in skin-draining lymph nodes six days post SC injection. **c, d**. Graphs showing % of Rho^+^ or Rho/DQ^+^ DC1 and DC2 in skin and skin-draining lymph nodes. **e**. Rho signals by DC1 and DC2 from human PBMCs after incubation with different dilutions of ACM-Rho.

To translate our results from mouse to human, we cultured primary human peripheral blood mononuclear cells (PBMCs) with ACM-Rhodamine in varying concentrations overnight and assessed their ability to take up ACM-Rhodamine by flow cytometry. We found that, compared to our mouse data, both human DC subsets, CD141^+^ DC1 and CD1c^+^ DC2, were capable of efficiently taking up ACM-Rhodamine as indicated by the high expression of Rhodamine in both populations, relative to the control (Fig. 2e).

In conclusion, both DC1 and DC2, are capable of efficiently taking up and processing ACMs and thus likely to present ACM-encapsulated antigen to naïve CD8^+^ T cells or CD4^+^ T cells. This may potentially induce a strong and durable adaptive immune response, which is highly desirable in the context of a vaccine.

### ACM-S1S2 + ACM-CpG formulation induced robust and durable neutralizing antibodies against SARS-CoV-2 in mice

Having established the DC-targeting property of ACM polymersomes, we proceeded to assess our ACM-spike vaccine formulations in C57BL/6 mice. Two doses of each formulation were administered at 2-week interval via subcutaneous injection and serum antibodies were examined on Day 13 (pre-boost) and Days 28, 40 and 54 (post-boost) (Fig. 3a). All antigens were injected at 5 μg per dose. Additionally, one group of mice received ACM-S1S2 + ACM-CpG formulation at 1/10^th^ dose (0.5 μg) for a limited dose-sparing investigation. Spike-specific IgG titers on Day 13 were moderate to low following a single dose of any formulation but increased dramatically by 21-255 folds on Day 28 after boost (Fig. 3b). Between the free and ACM-encapsulated antigen (S2, trimer or S1S2), a trend of higher IgG titer was observed in the latter, particularly after boost, suggesting that ACM technology enhanced the immunogenicity of each antigen. Between mice immunized with encapsulated trimer or S1S2 protein, Day 28 mean IgG titers were comparable at 1.0 × 10^5^ and 0.9 × 10^5^, respectively, suggesting similar immunogenicity. Focusing on our S1S2 protein, we saw progressive increase in Day 28 IgG titers with co-administration of CpG adjuvant, especially ACM-encapsulated CpG. The highest IgG response was achieved with the ACM-S1S2 + ACM-CpG formulation (mean titer of 8.5 × 10^5^), which even at 1/10^th^ dose elicited a robust IgG response (mean titer of 7.5 × 10^5^). To determine the durability of the IgG response, we continued monitoring mice up to Day 54. To the best of our knowledge, no other subunit vaccine developer had investigated antibody response in mice to such a late time point. A steady decrease in IgG titer was observed in each formulation (Fig. 3b), which resembled the decline after natural SARS-CoV-2 infection (32). Nevertheless, it was reported that viral neutralizing titers remained stable despite the decrease in IgG and hence we examined neutralizing responses next.

**Figure 3.**
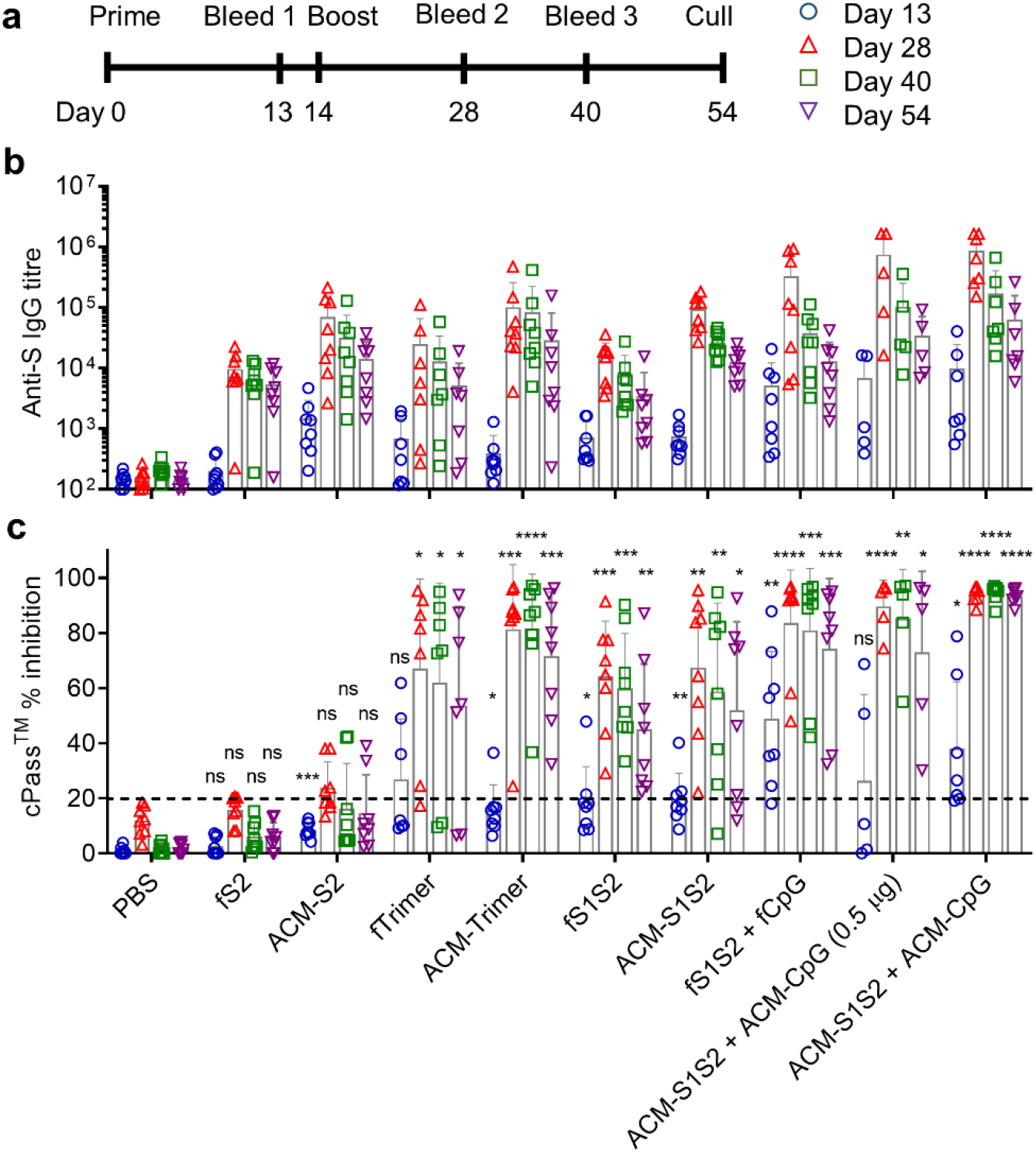
ACM-S1S2 + ACM-CpG vaccine elicited a vigorous SARS-CoV-2-specific antibody response. **a**. Immunization and blood collection schedule. C57BL/6 mice were subcutaneously immunized twice at 5 μg of antigen per dose (unless stated otherwise). **b**. Spike-specific total IgG. End point ELISA IgG titers were determined on plates coated with spike protein. **c**. Surrogate virus neutralization test. Neutralizing activity was determined using an ELISA-based cPass™ kit that assessed antibodies blocking the interaction between RBD and ACE2 receptor. A cut-off of 20% inhibition (horizontal dashed line) is used to identify seropositive samples. The different vaccine formulations being evaluated are indicated on the X-axis. Bar graphs depict means; error bars depict standard deviations. Statistical comparisons are made with respect to the PBS control at each time point using two-way ANOVA with Dunnett’s multiple comparison. *: P ≤ 0.05; **: P ≤ 0.01; ***: P ≤ 0.001; ****: P ≤ 0.0001; ns: not significant.

We adopted a multi-step approach to identify potentially neutralizing serum samples in a BSL-1/2 setting before doing a final validation against live virus in BSL-3. The first step involved the cPass™ kit, an FDA-approved, competitive ELISA-based assay that measured neutralizing antibodies blocking the interaction between recombinant RBD and ACE2 proteins. Crucially, this kit had been validated against patient sera and live SARS-CoV-2 and was shown to discriminate patients from healthy controls with 99.93% specificity and 95-100% sensitivity (33). Consistent with the low IgG titers on Day 13 (Fig. 3b), immune sera from different vaccine formulations generally showed little to no inhibition of RBD-ACE2 binding at 1:20 dilution (Fig. 3c), with the exception of the fS1S2 + fCpG and ACM-S1S2 + ACM-CpG mouse groups which exhibited seroconversion rates of 7/8 and 5/7, respectively. Next, we focused on sera collected after boost. Mice administered with free or ACM-encapsulated S2 protein continued showing little to no inhibitory activity from Day 28 to Day 54 (Fig. 3c), confirming the absence of neutralizing epitopes in S2. The spike trimer and S1S2 protein (free or encapsulated) generated highly variable responses on Day 28 that quickly declined at later time points. Strikingly, the ACM-S1S2 + ACM-CpG formulation elicited high levels of activity in all mice on Day 28 at 1:20 serum dilution, with an average inhibition of 94%. Moreover, levels of activity remained uniformly high till Day 54, indicating a durable response. To confirm these findings, we performed pseudovirus neutralization test on Day 28 sera from five key groups: ACM-S2, ACM-trimer, ACM-S1S2 and ACM-S1S2 + ACM-CpG (0.5 μg and 5 μg dosage groups). As expected, ACM-S2 failed to generate neutralizing antibodies against SARS-CoV-2 spike-pseudotyped virus (IC_50_ titer < 40; Fig. 4a). For the ACM-trimer and ACM-S1S2 mouse groups, partial seroconversion was observed with 7/8 and 4/8 mice, respectively, showing a positive response (IC_50_ titer ≥ 40). Finally, the ACM-S1S2 + ACM-CpG mouse group showed complete seroconversion with a mean IC_50_ titer of 789. Interestingly, even the 1/10^th^ (0.5 μg) dose remained highly efficacious, eliciting seroconversion in 5/5 mice with a mean titer of 773.

**Figure 4.**
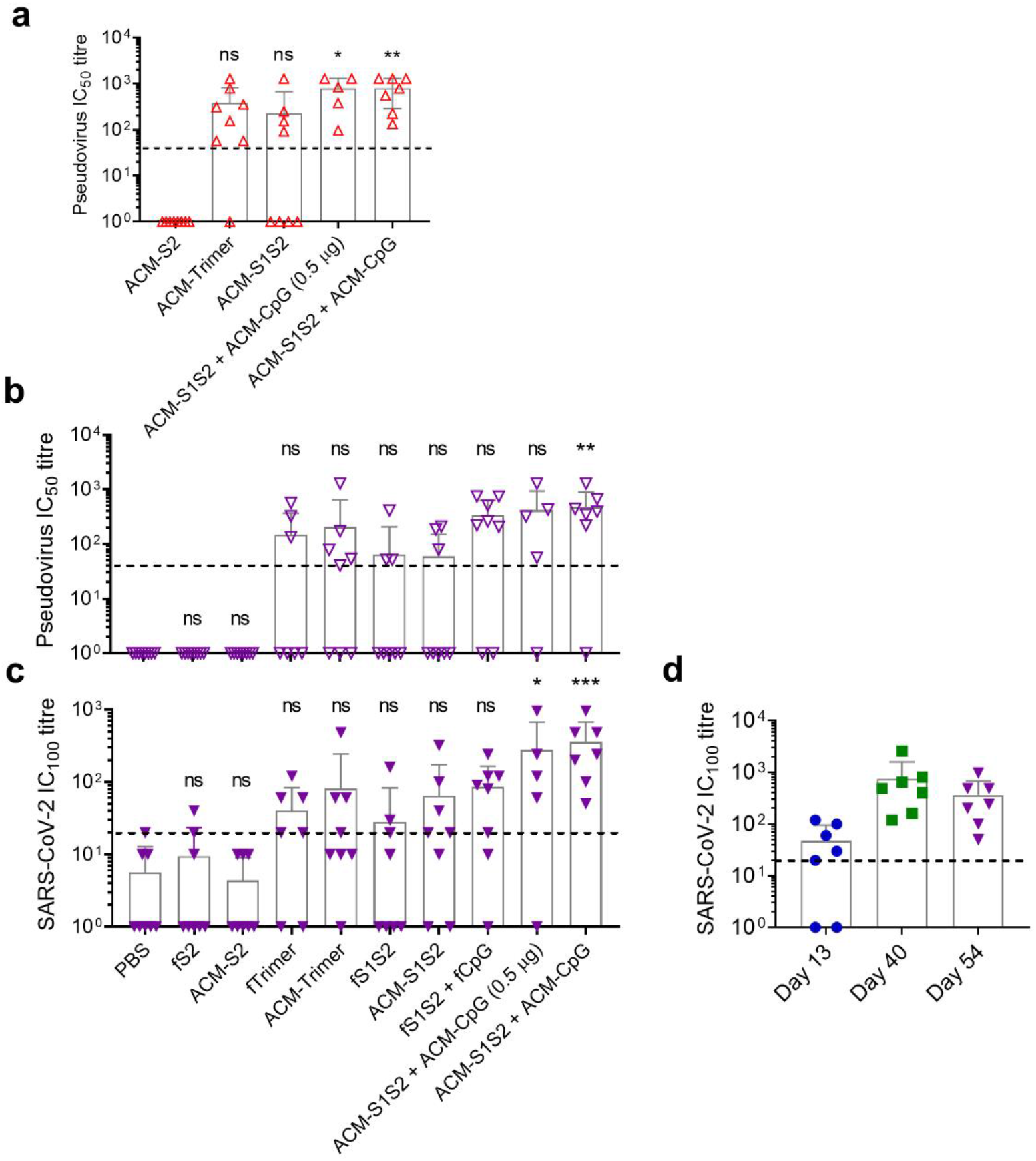
ACM-S1S2 + ACM-CpG vaccine elicited a robust and durable neutralizing antibody response. **a**. Day 28 sera from five key mouse groups were tested against SARS-CoV-2 spike-pseudotyped lentiviral particles to determine IC_50_ titres. **b**. IC_50_ neutralizing titers on Day 54 determined against SARS-CoV-2 spike-pseudotyped lentiviral particles. **c**. IC_100_ neutralizing titers on Day 54 determined against live SARS-CoV-2. Lower limits of detection are indicated by horizontal dashed lines; samples below threshold are assigned a nominal value of 1. The different vaccine formulations being evaluated are indicated on the X-axis. Bar graphs depict means; error bars depict standard deviations. Statistical comparisons are made with respect to the ACM-S2 or PBS group using ordinary one-way ANOVA with Dunnett’s multiple comparison. *: P ≤ 0.05; **: P ≤ 0.01; ***: P ≤ 0.001; ns: not significant. **d**. Kinetics of neutralizing titres from ACM-S1S2 + ACM-CpG-immunized mice.

We proceeded to analyse sera from the last time point (Day 54) by pseudovirus and live SARS-CoV-2 neutralization tests (Fig. 4b, c). Neutralizing responses across mouse groups were generally moderate to low, with many mice falling below respective limits of detection. Only the ACM-S1S2 + ACM-CpG group retained high neutralizing titers with a mean IC_50_ titer of 475 against pseudovirus (Fig. 4b), and IC_100_ titer of 359 against live SARS-CoV-2 (Fig. 4c). Even the 1/10^th^ dose demonstrated good efficacy, inducing mean IC_50_ titer of 416 against pseudovirus and IC_100_ titer of 276 against SARS-CoV-2. Between the two neutralizing assays, results were generally in strong agreement (Pearson correlation coefficient: 0.83; Supplementary Fig. 6) though some discrepancies were observed. In particular, we noticed one mouse from the ACM-S1S2 + ACM-CpG group that was seemingly seronegative on Day 54 by pseudovirus neutralization test but was consistently seropositive across time points by the cPass™ assay as well as live SARS-CoV-2 neutralization test. This discrepancy may arise from the relatively high threshold of 1:40 serum dilution of the pseudovirus assay, which was needed to address high background activity of some naïve mice that we and others (34) have noticed. At the same time, we must acknowledge that our understanding of the SARS-CoV-2 infection process is likely incomplete. The design of the pseudovirus virus assay is based on the interactions between SARS-CoV-2 spike protein and host cell ACE2 receptor and proteases (35, 36), as well as the ability of neutralizing antibodies to disrupt these interactions (37). It is possible antibodies may neutralize in a mechanism not recapitulated by this assay. Therefore, results should be validated with live SARS-CoV-2 neutralization test, which is the gold standard.

To better understand the kinetics of the neutralizing response after ACM-S1S2 + ACM-CpG vaccination, sera from Days 13 and 40 were also assessed by live virus neutralization test (Fig. 4d; Day 28 sera unavailable due to the earlier pseudovirus test). A single dose of ACM-S1S2 + ACM-CpG elicited partial seroconversion with a mean IC_100_ titer of 47 on Day 13, whereas two doses resulted in a sharp rise in IC_100_ titer to 737 on Day 40. Together with the earlier serum IgG data, this strongly supported a prime-boost regimen to induce robust neutralizing titers. Altogether, we demonstrated that ACM-S1S2 + ACM-CpG at 5 μg dose induced high levels of neutralizing antibodies in all mice. Moreover, neutralizing titers persisted at least 40 days after the last administration, suggesting a durable response.

### ACM-S1S2 + ACM-CpG formulation induced Th1-biased, functional memory T cells against SARS-CoV-2 spike protein in mice

To evaluate spike-specific T cell responses, splenocytes were harvested from all mice on Day 54 and stimulated *ex vivo* with an overlapping peptide pool covering the spike protein. T cell function was measured by intracellular cytokine staining. At this late time point (40 days after boost), activated T cells would have progressed beyond the initial expansion phase and entered contraction/memory phase (38). To the best of our knowledge, only one mRNA vaccine had been investigated for murine T cell responses at the late time point of seven weeks after boost (39). Memory-phenotype CD4^+^ and CD8^+^ T cells were identified by gating on the respective CD44^hi^ subpopulations (Supplementary Fig. 7a). Among the S1S2 vaccine groups, only the ACM-S1S2 + ACM-CpG formulation (5 or 0.5 μg dose) induced highly significant increase in IFNγ-, TNFα- or IL-2-expressing CD4^+^ T cells in response to spike peptide stimulation (Fig. 5a and Supplementary Fig. 7b. For the S2 and trimer mouse groups, no significant increase in Th1 cytokine-producing CD4^+^ T cells was detected above baseline (Supplementary Fig. 7d). With regards to Th2 cytokines, IL-4 was not detected in any mouse group whereas IL-5 was consistently elevated in non-adjuvanted S1S2-, S2- or trimer-immunized mice (Fig. 5a and Supplementary Fig. 7d, respectively), indicating a Th2-biased immune response. The Th2 skew was also evident from their IgG1:IgG2b ratios (Fig. 5c). Strikingly, production of IL-5 was strongly suppressed by co-administration of CpG. In particular, the ACM-S1S2 + ACM-CpG formulation (5 or 0.5 μg dose) produced a clear Th1-polarized profile, which was also reflected by an IgG1:IgG2b ratio < 1 (Fig. 5a & c, respectively). With regards to CD8^+^ T cells (Supplementary Fig. 7c), IFNγ was the predominant response in the ACM-S1S2 + ACM-CpG (5 μg dose) group, with all mice showing activity above baseline (Fig. 5b). In addition, some mice had slight expression of TNFα and IL-2 though the average frequencies of responding cells were not significantly elevated. A similar cytokine profile was seen in the ACM-S1S2 group, though only 5/8 mice had IFNγ responses above baseline. For the remaining mouse groups, CD8^+^ T cell responses were not significantly elevated (Fig. 5b and Supplementary Fig. 7e). Collectively, ACM-S1S2 + ACM-CpG (5 μg dose) induced in all mice functional memory CD4^+^ and CD8^+^ T cells that were readily detected even after 40 days from the last administration. Additionally, CD4^+^ T cells exhibited a Th1-skewed cytokine profile, which was also reflected in the predominance of IgG2b over IgG1.

**Figure 5.**
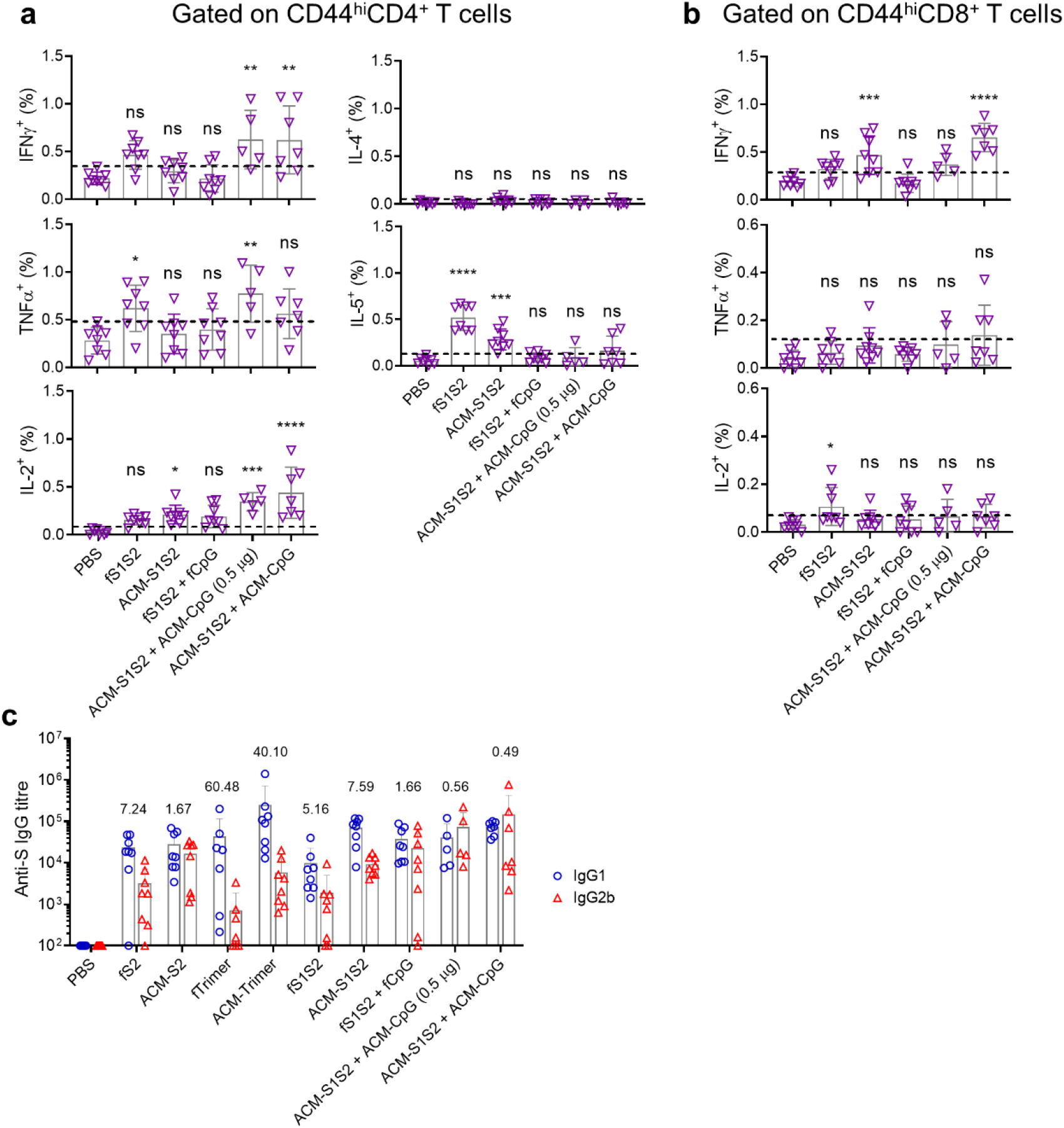
ACM-S1S2 + ACM-CpG vaccine elicited functional memory CD4^+^ and CD8^+^ T cells. Spleens were harvested on Day 54 (40 days after boost) and splenocytes (including those from PBS controls) were stimulated *ex vivo* with an overlapping peptide pool covering the SARS-CoV-2 spike protein. T cell responses were determined by intracellular cytokine staining. **a**. Th1 (IFNγ, TNFα and IL-2) and Th2 (IL-4 and IL-5) cytokine production by CD44^hi^CD4^+^ T cells. **b**. IFNγ, TNFα and IL-2 production by CD44^hi^CD8^+^ T cells. Baselines (horizontal dashed lines) are assigned according to PBS controls and readings above them are considered antigen-specific. The different formulations being evaluated are indicated on the X-axis. Bar graphs depict means; error bars depict standard deviations. Statistical comparisons are made with respect to the PBS control using ordinary one-way ANOVA with Dunnett’s multiple comparison. *: P ≤ 0.05; **: P ≤ 0.01; ***: P ≤ 0.001; ****: P ≤ 0.0001; ns: not significant. **c**. Spike-specific IgG1 and IgG2b titers of Day 54 sera. End point titers were determined on plates coated with spike protein. Average IgG1:IgG2b ratios are indicated above bar graphs.

In summary, ACM-S1S2 + ACM-CpG induced functional memory CD4^+^ and CD8^+^ T cells that could be detected 40 days after the last administration. The efficient uptake of ACM vesicles by cDC1 is likely important for generating CD8^+^ T cell immunity, given cDC1’s ability to efficiently cross-present (40). Accordingly, other groups have reported efficient delivery of antigens to DCs using nanoparticles, which resulted in cross-presentation and CD8^+^ T cell priming (41-43). In our hands, we have demonstrated spike-specific CD8^+^ T cell responses in mice vaccinated with ACM-S1S2 but not free S1S2 protein.

Inclusion of CpG in the vaccine formulation confers several benefits. It potently activates DCs to upregulate co-stimulatory molecules, including CD40, CD80 and CD86 (44), which promotes T cell activation and B cell antibody class switch and secretion (45, 46). Binding of CpG to TLR-9 triggers MAPK and NF-kB signalling that results in pro-inflammatory cytokine production and a Th1-skewed immune response (47, 48). In our study, such polarization is clearly demonstrated by the cytokine profile of CD4^+^ T cells and the IgG1:IgG2b ratio of the CpG-containing vaccine formulations. In the absence of CpG, we consistently observed IL-5 production which fits a broader picture of an inherent Th2 skew from immunizing with protein antigens of viral and non-viral origins (49, 50). From a safety standpoint, this represents a potential risk of Th2 immunopathology, best exemplified by whole-inactivated RSV vaccines (25, 51). Accordingly, such vaccines primed the immune system for a Th2-biased response during actual infection and the resultant production of Th2 cytokines promoted increased mucus production, eosinophil recruitment and airway hyperreactivity. Therefore, skewing of the immune response to Th1 by CpG is likely to improve vaccine safety.

We have shown that neutralizing titers can remain stable despite rapidly declining total IgG, which is consistent with SARS-CoV-2-infection in humans (32). This may be due to affinity maturation which progressively selects for high avidity, strongly neutralizing antibodies while excluding weaker binders. Additionally, compared to the neutralizing titers measured in convalescent patients recruited in Singapore (52), it appears that our vaccine formulation may be more efficient in triggering neutralizing antibodies. Although the role of antibodies in Covid-19 remains to be established, it is reasonable to regard neutralizing antibodies as a potential correlate of protection. Reports of asymptomatic or mild patients producing widely varying neutralizing antibody levels, including a minority with no detectable neutralizing response (4, 53), underscore the unpredictability of a natural infection. In this regard, our vaccine can perhaps facilitate the induction of a more uniform neutralizing antibody response.

The role of T cells in SARS-CoV-2 is arguably less clear than antibodies. Nevertheless, several studies have confirmed the induction of a T cell response following infection. Early in the adaptive immune response against SARS-CoV-2, T cells are robustly activated (54). Patients who recovered from SARS in 2003 possessed memory T cells that could be detected 17 years after (52). Additionally, individuals with no history of SARS, Covid-19 or contact with individuals who had SARS and/or Covid-19 possessed cross-reactive T cells that may be generated by a previous infection with other betacoronaviruses (52). These data suggested that the SARS-CoV-2-specific T cell response may be similarly durable. In a study examining the T cell specificities of Covid-19 convalescent patients, spike-specific CD4^+^ T cells were consistently detected whereas CD8^+^ T cells were present in most subjects (10). This implies that a spike-based vaccine may generate a cellular immune response that largely recapitulates the CD4^+^ T cell profile of a natural infection, albeit with a narrower CD8^+^ T cell repertoire.

One major challenge in creating a pandemic vaccine is generating sufficient doses of high-quality antigen to rapidly meet global demand. As such, dose-sparing strategies are critical, and this has traditionally been achieved using adjuvants. Based on our work, we believe that ACM technology together with an adjuvant can further augment the dose-sparing effect. We have shown this approach to greatly improve vaccine immunogenicity, such that even the 1/10^th^ dose retains a substantial level of efficacy. While further titration experiments are required to determine the optimum dose, the present limited investigation strongly supports the use of ACM technology to address limited antigen availability in a pandemic.

## Methods

Methods, including statement of data availability, statement of restriction on biological material availability, and GenBank accession number of reference SARS-CoV-2 genomic sequence, are available in the online version of this paper.

## Acknowledgements

We would like to thank Ting Yan Aw (ACM Biolabs), Jianyao Nicholas Chng (ACM Biolabs) and Liam Thomas Martin (ACM Biolabs) for assisting in protein expression in insect cells and optimization of protein purification; James Ho (CBSS, NTU) for helping with Cryo EM sample preparation and Andrew Wong (NTU) for Cryo EM sample measuring; Soh Yee Joey Poh and the staff of BRC (A*STAR) for assisting in mouse immunization and sample collection. We would like to acknowledge IAF-ICP, A*STAR grant for ACM-SIgN collaboration and NHIC Gap Funding Award (NHIC-COV19-2005008) for collaboration with Dr. Francesca Lim, Singapore General Hospital, and Dr. Danielle E. Anderson, Duke-NUS Medical School, Singapore. The work performed in NUS/NUHS was supported by NUHS Research Office under Project Number NUHSRO/2020/033/RO5+5/CORONAVIRUS/LOA (WBS R-571-000-071-733).

## Author contributions

J.H.L., A.K.K., T.A.C., R.J.D., T.W.C., W.W.W.Y., N.K.M.I., K.T.N. and D.E.A. designed and performed the experiments. Y.J.T., R.J.D., F.G. and M.N. designed the experiments, analyzed the data and provided critical intellectual input. J.H.L., A.K.K., S.V., R.J.D. and M.N. wrote the manuscript. All authors contributed to manuscript revisions and approved the final version.

## Additional information

Supplementary information is available in the online version of the paper.

## Competing interests

The authors declare the following competing financial interests: D.E.A. and Y.J.T. developed the cPass™ kit; J.H.L, A.K.K., T.A.C., T.W.C., W.W.W.Y., and M.N. are employees of ACM Biolabs Pte Ltd; F.G. is part of the ACM SAB.

The authors declare no non-financial competing interests.

## Methods

### Materials

Murine CpG 1826 (T*C*C*A*T*G*A*C*G*T*T*C*C*T*G*A*C*G*T*T, where * denotes phosphodiester backbone) was purchased from InvivoGen. Rhodamine B-terminated PEG_13_-*b*-PBD_22_ was purchased from Polymer Source Inc. DQ ovalbumin protein (OVA-DQ) was purchased from Life Technologies, Thermo Fisher Scientific. 1,2-dioleoyl-3-trimethylammonium-propane (DOTAP) was from Avanti Polar Lipids. Triton X-100 was from MP Biomedicals. All other chemicals were purchased from Sigma-Aldrich unless stated otherwise. The trimeric spike protein was purchased from ACROBiosystems and the S2 domain protein from Sino Biological.

### Protein expression

Recombinant SARS-CoV-2 spike protein containing only the ectodomain (hereby referred to as “S1S2”), from Genbank entry MN908947.3, with a mutated furin cleavage site (NSPRRAR → NSNQSAR) and a melittin secretion leader, was expressed via T.ni insect cells (Hi5, Thermo Fisher Scientific). The gene of interest was placed into the Bac-to-Bac system (Thermo Fisher Scientific), transfected and passaged in Sf9 cells (Thermo Fisher Scientific) until a high titre was achieved. T.ni cells, diluted to 1.5 × 10^6^ cells/ml, were infected at a MOI of 0.1 and left to incubate (27°C for 96 h, shaking at 125 rpm). The cell culture was harvested, and the cells removed by centrifugation (3,500 x g for 15 min at 4°C) and clarified by 0.22 µm filtration. The media containing the protein of interest was first concentrated to a tenth of the original volume via Tangential flow filtration hollow fibre cassettes (10 kDa Hollow fibre cassette; Cytiva), followed by 5 volumes worth of diafiltration into IEX binding buffer (20 mM Phosphate, 50 mM NaCl, 5% sucrose, 5% glycerol, 0.025% tween 20, 1 mM EDTA, pH 4.6). The protein was initially purified by first binding the sample in a HiTrap FF SP column (5 ml; Cytiva) using a GE AKTA system loaded with Unicorn software, set at 2 ml/min. Once the sample had been loaded and washed with 5 column volumes of IEX binding buffer, the protein of interest was eluted off the column by switching to IEX elution buffer (20 mM Phosphate, 50 mM NaCl, 5% sucrose, 5% glycerol, 0.025% tween 20, 1 mM EDTA, pH 7.6). The eluted sample was concentrated using a Vivaspin concentrator (10 kDa, 15 ml, PES; Sartorius) to a 5 ml volume. The protein was polished by loading 2.5 ml of sample in a 5 ml loading loop onto a Hiload 16/60 Superdex 200 Prep Grade column, running with SEC buffer (20 mM Phosphate, 150 mM NaCl, 5% sucrose, pH 7.6) at 1 ml/min. Purified protein was analysed for size by injection of 100 µl of sample into a Superdex 200 increase 10/300 GL column using a GE AKTA system running at 0.75 ml/min. Molecular mass of the protein was calculated via comparison with a Gel filtration calibration kit HMW (containing a mixture of Thyroglobulin, Ferritin, Aldose and Conalbumin; Cytiva).

### Preparation of ACM-antigen polymersomes

ACM polymersomes encapsulating spike trimer, S1S2 and S2 proteins were prepared by the solvent dispersion method, followed by extrusion. A 400 mg/ml stock solution of DOTAP and PEG_13_-*b*-PBD_22_ polymer were prepared by dissolving solid DOTAP and polymer in tetrahydrofuran (THF). 0.15 equivalents (1.5 µmol) of DOTAP stock solution and 0.85 equivalents (8.5 µmol) of polymer stock solution were mixed in a 2 ml glass vial and vortexed to prepare Solution A. After mixing, Solution A was aspirated in a 50 µl Hamilton glass syringe. A 1 ml solution of 100 µg/ml antigen was placed in a 5 ml glass test tube (Solution B). Solution A was added slowly to 1 ml of Solution B while constantly mixing (600-700 rpm) at room temperature. A turbid solution was obtained. The resultant solution was extruded 21 times through a 200 nm membrane filter (Avanti Polar Lipids) using a 1 ml mini-extruder (Avanti Polar Lipids) to get monodispersed ACM-antigen vesicles. Non-encapsulated antigens were removed by overnight dialysis. Encapsulation of antigen were quantified by densiometric analysis using a known BSA standards in Fiji ImageJ software (v. 1.52a).

### Preparation of ACM-CpG polymersomes

ACM-CpG polymersomes were prepared by the solvent dispersion method above, followed by extrusion. 50 µl of the 400 mg/ml stock solution containing DOTAP and PEG_13_-*b*-PBD_22_ polymer was added dropwise to 1 ml CpG solution. A turbid solution was obtained. The resultant solution was extruded 21 times through a 200-nm membrane filter using a 1 ml mini-extruder to get monodispersed ACM-CpG polymersomes. Unencapsulated CpG was removed by overnight dialysis using 300 kDa molecular weight cut-off (MWCO) regenerated cellulose membrane (Spectrum Laboratories Inc.) against PBS, pH 7.4 at 4°C.

### Preparation of ACM-Rhodamine and ACM-Rhodamine-OVA-DQ

ACM-Rhodamine and ACM-Rhodamine-OVA-DQ were prepared by the thin-film rehydration method, followed by extrusion. A 9.9 mg of PEG_13_-*b*-PBD_22_ polymer in chloroform were mixed with 0.1 mg Rhodamine B-terminated PEG_13_-*b*-PBD_22_ in chloroform with a ratio of 99:1 w/v shaken in a round bottom flask. After mixing, chloroform was removed by rotary evaporator followed by drying for 1 h at high vacuum. A 1 ml solution of 100 µg/ml OVA-DQ was placed in the flask for the preparation of ACM-Rhodamine-OVA-DQ; for ACM-Rhodamine, 1 mL buffer was added. The solution was stirred at 600-700 rpm for overnight at 4°C. A pink coloured turbid solution was obtained. The resultant solution was extruded 21 times through a 200-nm membrane filter (Avanti Polar Lipids) using a 1 mL mini-extruder (Avanti Polar Lipids) to get monodispersed ACM nanoparticles. Non-encapsulated OVA-DQ was removed by overnight dialysis against 1X PBS.

### Particle size measurement by dynamic light scattering (DLS)

DLS was performed on the Zetasizer Nano ZS system (Malvern Panalytical). 100 µl of the 20-fold diluted, purified, filtered sample was placed in a micro cuvette (Eppendorf^®^ UVette; Sigma-Aldrich) and an average of 30 runs (10 s per run) was collected using the 173° detector.

### Quantification of spike protein by SDS-PAGE

20 µl of ACM-spike protein or free spike protein at known concentrations was added to microcentrifuge tubes. 2 µl of 25% Triton X-100 was added to each sample and incubated for 30 min at 25°C to lyse ACM vesicles. Next, 20 µl of 1X gel loading dye buffer was added and tubes were shaken at 95°C for 10 min. 20 µl of each sample was migrated on 4-12% Bis-Tris SDS-PAGE gel at 140 V for 40 min. The completed gel was fixed and then stained with SYPRO^®^ Ruby protein gel stain (Molecular Probes, Thermo Fisher Scientific).

### Western blot

Proteins were transferred from SDS-PAGE gel to PVDF membrane using the iBlot 2 Dry Blotting System (Thermo Fisher Scientific). The membrane was blocked 1 h at room temperature with 5% w/v non-fat milk dissolved in TBST (Tris-buffered saline with 0.1% v/v Tween-20). Mouse serum raised against a recombinant SARS-CoV-2 spike protein (purchased from Sino Biological) was diluted 1:6,000 and incubated with the membrane for 1 h at room temperature. The membrane was washed thrice with TBST for a total of 30 min before incubating 1 h at room temperature with HRP-conjugated goat anti-mouse secondary antibody at a 1:10,000 dilution. After three final washes with TBST, the membrane was briefly incubated with ECL substrate (Pierce, Thermo Fisher Scientific). Chemiluminescent signals were captured using the ImageQuant LAS 500 system (Cytiva).

### Quantification of CpG by fluorescence

20 µl of ACM-CpG or free CpG at known concentrations were added to a 384-well black plate. 20 µl of PBS with 10% Triton X-100 was added into each well, and the plate was incubated for 30 min at 25°C to lyse ACM vesicles before adding 10 µl of 20X SYBR™ Safe DNA gel stain (Invitrogen, Thermo Fisher Scientific). The plate was incubated for 5 min at 25°C and fluorescence was measured (excitation – 500 nm; emission – 530 nm) using a plate reader (Biotek).

### Cryogenic-transmission electron microscopy (Cryo-TEM)

For cryo-TEM, 4 µL of the samples containing ACM-S1S2, ACM-CpG, and ACM-S1S2 + ACM-CpG vesicles (5 mg/ml) were adsorbed onto a lacey holey carbon-coated Cu grid, 200 mesh size (Electron Microscopy Sciences). The grid was surface treated for 20 s via glow discharge before use. After surface treatment, 4 µl sample was added and the grid was blotted with Whatman filter paper (GE Healthcare Bio-Sciences) for 2 s with blot force 1, and then plunged into liquid ethane at -178 °C using Vitrobot (FEI Company). The cryo-grids were imaged using a FEG 200 keV transmission electron microscope (Arctica; FEI Company) equipped with a direct electron detector (Falcon II; Fei Company). Images were analyzed in Fiji ImageJ software (v. 1.52a) and membrane thickness of vesicles were calculated by counting at least 20 particles.

### Mice (investigation of DC targeting by ACM polymersomes)

C57BL/6 mice were purchased from InVivos. All mice were maintained in the Singapore Immunology Network (SIgN) animal facility before use at 7-10 weeks of age. All experiments and procedures were approved by the Institutional Animal Care and Use Committee of the Biological Resource Center (Agency for Science, Technology and Research, Singapore) in accordance with the guidelines of the Agri-Food and Veterinary Authority and the National Advisory Committee for Laboratory Animal Research of Singapore (ICUAC No. 181357).

### Mice (vaccination)

This study was performed at the Biological Resource Center (Agency for Science, Technology and Research, Singapore). Female C57BL/6 mice were purchased from InVivos and used at 8-9 weeks of age. Seven to eight mice were assigned to each vaccine formulation, unless stated otherwise. Mice were administered 5 μg of a respective antigen (free or encapsulated) with or without 5 μg CpG adjuvant (free or encapsulated) in 200 μl volume per dose via the subcutaneous route, for one prime and one boost separated by 14 days. Blood was collected on days 13, 28, 40 and 54; spleens were collected on the final time point of day 54. The study was done in accordance with approved IACUC protocol 181137.

### Mouse tissue preparation and data analysis for flow cytometry

Mice were injected subcutaneously with 100 μl PBS, 100 μl ACM-Rhodamine or 100 μl ACM-Rhodamine-OVA-DQ and analysed on day 1, 3 or 6 post injection. Back skin from the injection site was harvested and placed in RPMI1640 (Gibco, Thermo Fisher Scientific) containing Dispase for 90 min at 37°C. The back skin and skin-draining LNs (separately) then were transferred into RPMI1640 containing DNaseI (Roche) and collagenase (Sigma-Aldrich), disrupted using scissors or tweezers, and digested for 30 min at 37 °C. Digest was stopped by adding PBS + 10 mM EDTA and cell suspensions were transferred into a fresh tube over a 70 μm nylon mesh sieve. If necessary, red blood cells were lysed using RBC lysis buffer (eBioscience™), and single cell suspensions were passed through a 70 μm nylon mesh sieve before further use. Single cell suspensions then were stained for flow cytometry analysis following standard protocols. Monoclonal antibodies against Ly6C (clone HK1.4), CD11b (clone M1/70), EpCAM (clone G8.8), CD64 (clone X54-5/7.1), and F4/80 (clone BM8) were purchased from BioLegend, CD11c (clone N418), CD103 (clone 2E7), CD8a (clone 53-6.7), and MHC-II (clone M5/114.15.2) were purchased from eBioscience, CD24 (clone M1/69), CD3 (clone 500A2), CD45 (clone 30-F11), CD49b (clone HMa2), and Ly6G (clone 1A8) were purchased from BD Bioscience, CD19 (clone 1D3) and Streptavidin for conjugation of biotinylated antibodies were purchased from BD Horizon. DAPI staining was used to allow identification of cell doublets and dead cells. Flow cytometry acquisition was performed on a 5-laser LSR II (BD) using FACSDiva software, and data subsequently analyzed with FlowJo v.10.5.3 (Tree Star).

### Intracellular cytokine staining

Single-cell suspensions of splenocytes were generated by pushing each spleen through a 70 μm cell strainer. Red blood cells were lysed using 1X RBC Lysis Buffer (eBioscience, Thermo Fisher Scientific) for 5 min at room temperature. Splenocytes were resuspended in complete cell culture medium (RPMI 1640 supplemented with 10% v/v heat-inactivated FBS, 50 µM β-mercaptoethanol, 2 mM L-glutamax, 10 mM HEPES and 100 U/ml Pen/Strep; all materials purchased from Gibco, Thermo Fisher Scientific) and seeded in a 96-well U-bottom plate at a density of ∼3 million per well. Splenocytes were incubated with an overlapping peptide pool covering the spike protein (JPT product PM-WCPV-S-1 Vials 1 and 2) along with functional anti-mouse CD28 and CD49d antibodies overnight at 37°C, 5% CO_2_. Peptides and antibodies were used at 1 μg/ml, respectively. Negative control wells were generated by incubating splenocytes with culture medium and costimulatory antibodies. Positive control wells were generated by incubating splenocytes with 20 ng/ml PMA (Sigma-Aldrich) and 1 μg/ml ionomycin (Sigma-Aldrich). The following morning, cytokine secretion was blocked with 1× brefeldin A (eBioscience) and 1× monensin (eBioscience) for 6 h. Subsequently, cells were stained with Fixable Viability Dye eFluor™ 455UV (eBioscience) at 1:1000 in PBS for 30 min at 4°C. Cells were washed with FACS buffer (1× PBS supplemented with 2% v/v heat-inactivated FBS and 1 mM EDTA) and stained for surface markers with the following antibodies purchased from BioLegend, eBioscience and BD: BUV395-CD45 (30-F11), Brilliant Violet 785™-CD3 (17A2), Alexa Fluor 700-CD4 (GK1.5), APC-eFluor 780-CD8 (53-6.7) and PE/Dazzle™ 594-CD44 (IM7). Antibodies were diluted 1:200 with FACS buffer and incubated with cells for 30 min at 4°C. Fixation and permeabilization was done using the Cytofix/Cytoperm™ kit (BD), according to manufacturer’s instructions. Intracellular cytokines were stained with the following antibodies: Alexa Fluor 488-IFNγ (XMG1.2), Brilliant Violet 650-TNFα (MP6-XT22), APC-IL-2 (JES6-5H4), PerCP-eFluor 710-IL-4 (11B11) and PE-IL-5 (TRFK5). Antibodies were diluted 1:200 with 1× Permeabilization Buffer and incubated with cells for 30 min at 4°C. Cells were washed with 1× Permeabilization Buffer and then resuspended in FACS buffer for analysis with the LSR II flow cytometer (BD). Approximately 600,000 total events were recorded for each sample. Data analysis was performed using FlowJo V10.6.2 software. Percentage of cytokine-positive events for immunized mouse groups were compared against PBS-control group. Responses above the background of the PBS-control group were considered spike-specific.

### ACM uptake in human PBMC

Blood samples were obtained from healthy donors after provided written informed consent to participate in research protocols approved by the Institutional Review Board of Singapore Immunology Network (SIgN). Peripheral blood mononuclear cells (PBMCs) were isolated by Ficoll-Paque (GE Healthcare) density gradient centrifugation of apheresis. Isolated PBMCs were cultured overnight in RPMI1640 medium supplemented with 2% human AB serum, 1% penicillin/streptomycin (Gibco, Thermo Fisher Scientific) in 96-well round-bottom tissue plates, at 37°C, with or without ACM-Rhodamine (1:50 or 1:250 dilution). PBMCs then were stained for flow cytometry following standard protocols, briefly, 5 × 10^6^ cells/tube were washed and incubated with live/dead blue dye (Invitrogen) for 30 min at 4 °C in PBS, followed by incubation in 5% heat-inactivated FCS for 15 min at 4°C (Sigma Aldrich). Antibodies were diluted in PBS with 2% FCS and 2 mM EDTA, added to the cells and incubated for 30 min at 4°C. PBMCs were stained with mouse anti-human monoclonal surface antibodies (mAbs) against CD45 (clone HI30), CD3 (clone SP34-2), CD19 (clone SJ25C1), CD20 (clone 2H7), CD14 (clone M5E2), and CD123 (clone 7G3) purchased from BD Biosciences, mAbs against HLA-DR (clone L243), CD16 (clone 3G8), and CD1c (clone L161) purchased from BioLegend and mAb against CD141 (clone AD5-14H12) purchased from Miltenyi. Flow cytometry was performed on a BD FACSFortessa (BD Biosciences). Data were analyzed using FlowJo v.10.5.3 (Tree Star).

### ACE2 binding assay

Spike protein was coated onto 96-well EIA/RIA high binding plate (Corning) in carbonate-bicarbonate buffer (15 mmol/L Na_2_CO_3_, 35 mmol/L NaHCO_3_; pH 9.6) at 200 ng per well, overnight at 4 °C. Plates were blocked with 2% BSA in TBS + 0.05% v/v Tween-20 for 1.5 h at 37°C. Three-fold serial dilutions of recombinant hACE2-Fc protein (12,000 ng/ml to 0.61 ng/ml; GenScript) were prepared in TBS buffer containing 0.5% w/v BSA and applied to the plate for 1 h at 37 °C. HRP-conjugated goat anti-human IgG (Fc specific; Sigma Aldrich) was diluted 1:10,000 and applied to the plate for 1 h at 37°C. ACE2 binding was visualized by addition of TMB substrate (Sigma-Aldrich) for 15 min at room temperature and the reaction was terminated with Stop Solution (Invitrogen, Thermo Fisher Scientific). Absorbance was measured at 450 nm using a microplate reader (Biotek). Background absorbance was subtracted and the EC_50_ value of the titration curve was determined using GraphPad Prism version 8.4.3 with five-parameter non-linear regression.

### SARS-CoV-2 spike-specific serum IgG

Homemade spike protein was coated onto 96-well EIA/RIA high binding plate (Corning) at 100 ng per well in PBS overnight at 4°C. Plates were blocked with 2% w/v BSA in PBS + 0.1% v/v Tween-20 for 1.5 h at 37°C. Mouse sera were serially diluted from an initial of 1:100 with blocking buffer and applied to the plate for 1 h at 37°C. HRP-conjugated goat anti-mouse IgG (H/L), anti-mouse IgG1 or anti-mouse IgG2b (each purchased from BioRad) was diluted in blocking buffer at 1:10,000, 1:4,000 and 1:4,000, respectively, and applied to the plate for 1 h at 37°C. Antibody binding was visualized by addition of TMB substrate for 10 min at room temperature and the reaction was terminated with Stop Solution. Absorbance was measured at 450 nm. Each titration curve was analysed via five-parameter non-linear regression (GraphPad Prism V8.4.3) to calculate endpoint titer, which was defined as the highest dilution producing an absorbance three times the plate background.

### Serum neutralizing antibody by competitive ELISA

The cPass™ SARS-CoV-2 Surrogate Virus Neutralization Test Kit (GenScript) was used according to manufacturer’s instructions. Briefly, each serum sample was diluted 1:10 using Sample Dilution Buffer and incubated with an equal volume of HRP-RBD solution for 30 min at 37°C. The mix was then applied to 8-well strips pre-coated with ACE2 protein for 15 min at 37°C. RBD-ACE2 binding was visualized by addition of TMB substrate for 15 min at room temperature. Reaction was terminated using Stop Solution and absorbance was measured at 450 nm. Inhibition of RBD-ACE2 binding was calculated using the formula: 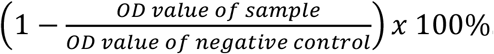.

### Pseudovirus neutralization test

Pseudotyped lentiviral particles harbouring the SARS-CoV-2 spike glycoprotein (S-pp) were generated by co-transfection of 293FT cells with S expression plasmid and envelope-defective pNL4-3.Luc.R-E-luciferase reporter vector. The S expression plasmid was constructed by cloning the codon-optimised spike gene (according to GenBank accession QHD43416.1) containing a 19 amino acid C-terminal truncation to enhance pseudotyping efficiency (36) into the pTT5 mammalian expression vector (pTT5LnX-coV-SP, a kind gift from Brendon John Hanson, Biological Defence Program, DSO National Laboratories, Singapore). The viral supernatant was collected 48-72 hours post-transfection, clarified by centrifugation, and stored at -80°C until use. S-pp titer was determined using a lentivirus-associated p24 ELISA kit (Cell Biolabs, Inc., San Diego, CA). CHO cells stably overexpressing human ACE2 (CHO-ACE2) (55) were seeded in 96-well plates 24 hour before transduction. Mouse serum samples were diluted 1:20 in culture medium, inactivated at 56°C for 30 min and sterilised using Ultrafree-MC centrifugal filters (Millipore, Burlington, MA). For S-pp neutralization assays, the serum samples were two-fold serially diluted six times and incubated with S-pp for 1 h at room temperature before the mixture was added to target cells in triplicate wells. Cells were incubated at 37°C for 48 h before being tested for luciferase activity using Bright-Glo™ Luciferase Assay System (Promega, Madison, WI). Luminescence was measured using a plate reader (Tecan Infinite M200) and after subtraction of background luminescence, the data were expressed as a percentage of the reading obtained in the absence of serum (cells + S-pp only), which was set at 100%. Dose-response curves were plotted with a four-parameter non-linear regression using GraphPad Prism 8 and neutralizing titers were reported as the serum dilution that blocked 50% S-pp entry (IC_50_). Samples that did not achieve 50% neutralization at the input serum dilution (1:40) were expressed as 1 while the neutralizing titer of samples that achieved more than 50% neutralization at the highest serum dilution (1:1280) were reported as 1280.

### SARS-CoV-2 neutralization test

Serum samples were serially diluted two-fold in DMEM supplemented with 5% v/v FBS, from an initial of 1:10 and incubated with equal volume of viral suspension (1 × 10^4^ TCID_50_/ml) for 90 min at 37°C. The mixture was transferred to Vero-E6 cells and incubated for 1 h at 37°C. The inoculum was removed, and cells were washed once with DMEM. Fresh culture medium was added, and cells were incubated for 4 days at 37°C. Assay was performed in duplicate. Neutralization titer was defined as the highest serum dilution that fully inhibited cytopathic effect (CPE).

## Data availability statement

The data that support the findings of this study are available from the corresponding author upon reasonable request.

## Restriction on availability of biological material statement

ACM polymersomes and ACM Covid-19 vaccine are proprietary materials of ACM Biolabs Pte Ltd and cannot be freely distributed upon request.

## References

1. Zhang R, Li Y, Zhang AL, Wang Y, Molina MJ. Identifying airborne transmission as the dominant route for the spread of COVID-19. Proceedings of the National Academy of Sciences. 2020;117(26):14857.

2. Sia SF, Yan L-M, Chin AWH, Fung K, Choy K-T, Wong AYL, et al. Pathogenesis and transmission of SARS-CoV-2 in golden hamsters. Nature. 2020;583(7818):834–8.

3. Richard M, Kok A, de Meulder D, Bestebroer TM, Lamers MM, Okba NMA, et al. SARS-CoV-2 is transmitted via contact and via the air between ferrets. Nature Communications. 2020;11(1):3496.

4. Wu Z, McGoogan JM. Characteristics of and Important Lessons From the Coronavirus Disease 2019 (COVID-19) Outbreak in China: Summary of a Report of 72 314 Cases From the Chinese Center for Disease Control and Prevention. JAMA. 2020;323(13):1239–42.

5. Gorbalenya AE, Baker SC, Baric RS, de Groot RJ, Drosten C, Gulyaeva AA, et al. The species Severe acute respiratory syndrome-related coronavirus: classifying 2019-nCoV and naming it SARS-CoV-2. Nature Microbiology. 2020;5(4):536–44.

6. Ke Z, Oton J, Qu K, Cortese M, Zila V, McKeane L, et al. Structures and distributions of SARS-CoV-2 spike proteins on intact virions. Nature. 2020.

7. Walls AC, Park Y-J, Tortorici MA, Wall A, McGuire AT, Veesler D. Structure, Function, and Antigenicity of the SARS-CoV-2 Spike Glycoprotein. Cell. 2020;181(2):281-92.e6.

8. Chi X, Yan R, Zhang J, Zhang G, Zhang Y, Hao M, et al. A neutralizing human antibody binds to the N-terminal domain of the Spike protein of SARS-CoV-2. Science. 2020;369(6504):650.

9. Rogers TF, Zhao F, Huang D, Beutler N, Burns A, He W-t, et al. Isolation of potent SARS-CoV-2 neutralizing antibodies and protection from disease in a small animal model. Science. 2020;369(6506):956.

10. Grifoni A, Weiskopf D, Ramirez SI, Mateus J, Dan JM, Moderbacher CR, et al. Targets of T Cell Responses to SARS-CoV-2 Coronavirus in Humans with COVID-19 Disease and Unexposed Individuals. Cell. 2020;181(7):1489-501.e15.

11. Krammer F. SARS-CoV-2 vaccines in development. Nature. 2020;586(7830):516–27.

12. Yadvinder SA, Dinesh SBaSKM. Adenoviral Vector Immunity: Its Implications and Circumvention Strategies. Current Gene Therapy. 2011;11(4):307–20.

13. Weng Y, Li C, Yang T, Hu B, Zhang M, Guo S, et al. The challenge and prospect of mRNA therapeutics landscape. Biotechnology Advances. 2020;40:107534.

14. Iqbal S, Blenner M, Alexander-Bryant A, Larsen J. Polymersomes for Therapeutic Delivery of Protein and Nucleic Acid Macromolecules: From Design to Therapeutic Applications. Biomacromolecules. 2020;21(4):1327–50.

15. Halperin A. Polymeric vs. Monomeric Amphiphiles: Design Parameters. Journal of Macromolecular Science, Part C. 2006;46(2):173–214.

16. Discher BM, Won Y-Y, Ege DS, Lee JCM, Bates FS, Discher DE, et al. Polymersomes: Tough Vesicles Made from Diblock Copolymers. Science. 1999;284(5417):1143.

17. Rideau E, Dimova R, Schwille P, Wurm FR, Landfester K. Liposomes and polymersomes: a comparative review towards cell mimicking. Chemical Society Reviews. 2018;47(23):8572–610.

18. Moon JJ, Suh H, Bershteyn A, Stephan MT, Liu H, Huang B, et al. Interbilayer-crosslinked multilamellar vesicles as synthetic vaccines for potent humoral and cellular immune responses. Nature Materials. 2011;10(3):243–51.

19. Scott EA, Stano A, Gillard M, Maio-Liu AC, Swartz MA, Hubbell JA. Dendritic cell activation and T cell priming with adjuvant- and antigen-loaded oxidation-sensitive polymersomes. Biomaterials. 2012;33(26):6211–9.

20. Stano A, Scott EA, Dane KY, Swartz MA, Hubbell JA. Tunable T cell immunity towards a protein antigen using polymersomes vs. solid-core nanoparticles. Biomaterials. 2013;34(17):4339–46.

21. Matoori S, Leroux J-C. Twenty-five years of polymersomes: lost in translation? Materials Horizons. 2020;7(5):1297–309.

22. Khan AK, Ho JCS, Roy S, Liedberg B, Nallani M. Facile Mixing of Phospholipids Promotes Self-Assembly of Low-Molecular-Weight Biodegradable Block Co-Polymers into Functional Vesicular Architectures. Polymers. 2020;12(4).

23. Ou X, Liu Y, Lei X, Li P, Mi D, Ren L, et al. Characterization of spike glycoprotein of SARS-CoV-2 on virus entry and its immune cross-reactivity with SARS-CoV. Nature Communications. 2020;11(1):1620.

24. Poh CM, Carissimo G, Wang B, Amrun SN, Lee CY-P, Chee RS-L, et al. Two linear epitopes on the SARS-CoV-2 spike protein that elicit neutralising antibodies in COVID-19 patients. Nature Communications. 2020;11(1):2806.

25. Graham BS. Rapid COVID-19 vaccine development. Science. 2020;368(6494):945.

26. de Alwis R, Chen S, Gan ES, Ooi EE. Impact of immune enhancement on Covid-19 polyclonal hyperimmune globulin therapy and vaccine development. EBioMedicine. 2020;55:102768-.

27. Fulginiti VA, Eller JJ, Downie AW, Kempe CH. Altered Reactivity to Measles Virus: Atypical Measles in Children Previously Immunized With Inactivated Measles Virus Vaccines. JAMA. 1967;202(12):1075–80.

28. Kim HW, Canchola JG, Brandt CD, Pyles G, Chanock RM, Jensen K, et al. RESPIRATORY SYNCYTIAL VIRUS DISEASE IN INFANTS DESPITE PRIOR ADMINISTRATION OF ANTIGENIC INACTIVATED VACCINE12. American Journal of Epidemiology. 1969;89(4):422–34.

29. Dress RJ, Wong AYW, Ginhoux F. Homeostatic control of dendritic cell numbers and differentiation. Immunology & Cell Biology. 2018;96(5):463–76.

30. Anderson DA, Dutertre C-A, Ginhoux F, Murphy KM. Genetic models of human and mouse dendritic cell development and function. Nature Reviews Immunology. 2020.

31. Merad M, Sathe P, Helft J, Miller J, Mortha A. The Dendritic Cell Lineage: Ontogeny and Function of Dendritic Cells and Their Subsets in the Steady State and the Inflamed Setting. Annual Review of Immunology. 2013;31(1):563–604.

32. Long Q-X, Tang X-J, Shi Q-L, Li Q, Deng H-J, Yuan J, et al. Clinical and immunological assessment of asymptomatic SARS-CoV-2 infections. Nature Medicine. 2020;26(8):1200–4.

33. Tan CW, Chia WN, Qin X, Liu P, Chen MIC, Tiu C, et al. A SARS-CoV-2 surrogate virus neutralization test based on antibody-mediated blockage of ACE2–spike protein– protein interaction. Nature Biotechnology. 2020;38(9):1073–8.

34. Nie J, Li Q, Wu J, Zhao C, Hao H, Liu H, et al. Establishment and validation of a pseudovirus neutralization assay for SARS-CoV-2. Emerging Microbes & Infections. 2020;9(1):680–6.

35. Hoffmann M, Kleine-Weber H, Schroeder S, Krüger N, Herrler T, Erichsen S, et al. SARS-CoV-2 Cell Entry Depends on ACE2 and TMPRSS2 and Is Blocked by a Clinically Proven Protease Inhibitor. Cell. 2020;181(2):271-80.e8.

36. Johnson MC, Lyddon TD, Suarez R, Salcedo B, LePique M, Graham M, et al. Optimized Pseudotyping Conditions for the SARS-COV-2 Spike Glycoprotein. Journal of Virology. 2020;94(21):e01062–20.

37. Jiang S, Hillyer C, Du L. Neutralizing Antibodies against SARS-CoV-2 and Other Human Coronaviruses. Trends in immunology. 2020;41(5):355–9.

38. Kim C, Fang F, Weyand CM, Goronzy JJ. The life cycle of a T cell after vaccination - where does immune ageing strike? Clinical and experimental immunology. 2017;187(1):71–81.

39. Corbett KS, Edwards D, Leist SR, Abiona OM, Boyoglu-Barnum S, Gillespie RA, et al. SARS-CoV-2 mRNA Vaccine Development Enabled by Prototype Pathogen Preparedness. bioRxiv. 2020:2020.06.11.145920.

40. Dudziak D, Kamphorst AO, Heidkamp GF, Buchholz VR, Trumpfheller C, Yamazaki S, et al. Differential Antigen Processing by Dendritic Cell Subsets in Vivo. Science. 2007;315(5808):107.

41. de Titta A, Ballester M, Julier Z, Nembrini C, Jeanbart L, van der Vlies AJ, et al. Nanoparticle conjugation of CpG enhances adjuvancy for cellular immunity and memory recall at low dose. Proceedings of the National Academy of Sciences. 2013;110(49):19902.

42. Kuai R, Ochyl LJ, Bahjat KS, Schwendeman A, Moon JJ. Designer vaccine nanodiscs for personalized cancer immunotherapy. Nature materials. 2017;16(4):489–96.

43. Luo M, Wang H, Wang Z, Cai H, Lu Z, Li Y, et al. A STING-activating nanovaccine for cancer immunotherapy. Nature Nanotechnology. 2017;12(7):648–54.

44. Iwasaki A, Medzhitov R. Toll-like receptor control of the adaptive immune responses. Nature Immunology. 2004;5(10):987–95.

45. Ma DY, Clark EA. The role of CD40 and CD154/CD40L in dendritic cells. The Many Faces of CD40: Multiple Roles in Normal Immunity and Disease. 2009;21(5):265–72.

46. Hubo M, Trinschek B, Kryczanowsky F, Tüttenberg A, Steinbrink K, Jonuleit H. Costimulatory Molecules on Immunogenic Versus Tolerogenic Human Dendritic Cells. Frontiers in Immunology. 2013;4:82.

47. Dalod M, Chelbi R, Malissen B, Lawrence T. Dendritic cell maturation: functional specialization through signaling specificity and transcriptional programming. The EMBO Journal. 2014;33(10):1104–16.

48. Weeratna RD, Brazolot Millan CL, McCluskie MJ, Davis HL. CpG ODN can redirect the Th bias of established Th2 immune responses in adult and young mice. FEMS Immunology & Medical Microbiology. 2001;32(1):65–71.

49. Balkovic ES, Florack JA, Six HR. Immunoglobulin G subclass antibody responses of mice to influenza virus antigens given in different forms. Antiviral Research. 1987;8(3):151–60.

50. Coutelier JP, van der Logt JT, Heessen FW, Warnier G, Van Snick J. IgG2a restriction of murine antibodies elicited by viral infections. Journal of Experimental Medicine. 1987;165(1):64–9.

51. Knudson CJ, Hartwig SM, Meyerholz DK, Varga SM. RSV Vaccine-Enhanced Disease Is Orchestrated by the Combined Actions of Distinct CD4 T Cell Subsets. PLOS Pathogens. 2015;11(3):e1004757.

52. Le Bert N, Tan AT, Kunasegaran K, Tham CYL, Hafezi M, Chia A, et al. SARS-CoV-2-specific T cell immunity in cases of COVID-19 and SARS, and uninfected controls. Nature. 2020;584(7821):457–62.

53. Ko J-H, Joo E-J, Park S-J, Baek JY, Kim WD, Jee J, et al. Neutralizing Antibody Production in Asymptomatic and Mild COVID-19 Patients, in Comparison with Pneumonic COVID-19 Patients. Journal of Clinical Medicine. 2020;9(7).

54. Sekine T, Perez-Potti A, Rivera-Ballesteros O, Strålin K, Gorin J-B, Olsson A, et al. Robust T Cell Immunity in Convalescent Individuals with Asymptomatic or Mild COVID-19. Cell. 2020;183(1):158-68.e14.

55. Lip K-M, Shen S, Yang X, Keng C-T, Zhang A, Oh H-LJ, et al. Monoclonal Antibodies Targeting the HR2 Domain and the Region Immediately Upstream of the HR2 of the S Protein Neutralize In Vitro Infection of Severe Acute Respiratory Syndrome Coronavirus. Journal of Virology. 2006;80(2):941.

