## Supplementary Figures for "Next generation vaccine platform: polymersomes as stable nanocarriers for a highly immunogenic and durable SARS-CoV-2 spike protein subunit vaccine"

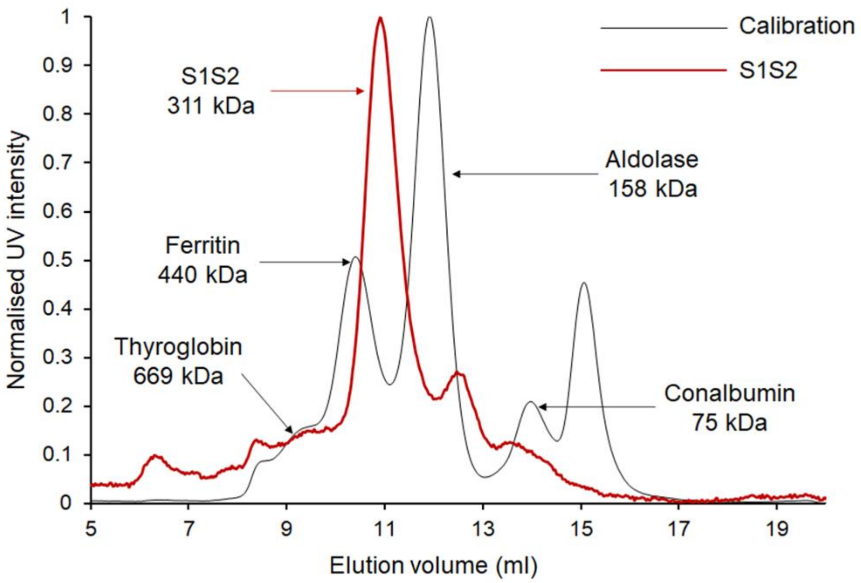

**Supplementary Figure 1 | Characterization of S1S2 protein by size exclusion chromatography.** Black trace: calibration curve. Red trace: purified S1S2 protein.

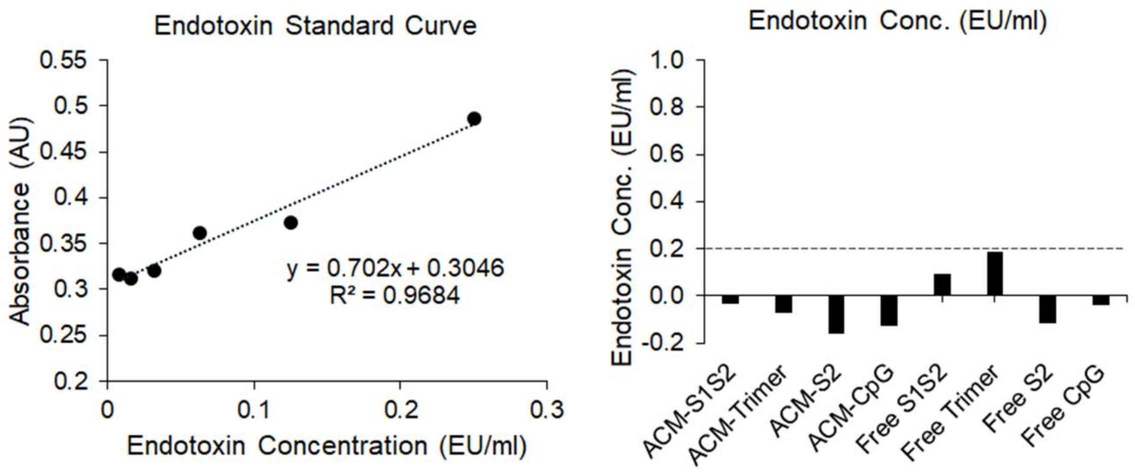

**Supplementary Figure 2 | Endotoxin measurement of ACM formulation.** Colorimetric HEK Blue cell-based endotoxin detection assay from InvivoGen showed negative endotoxic level for all ACM formulation and below 0.2 EU/ml endotoxin level for free S1S2 protein and free trimer protein.

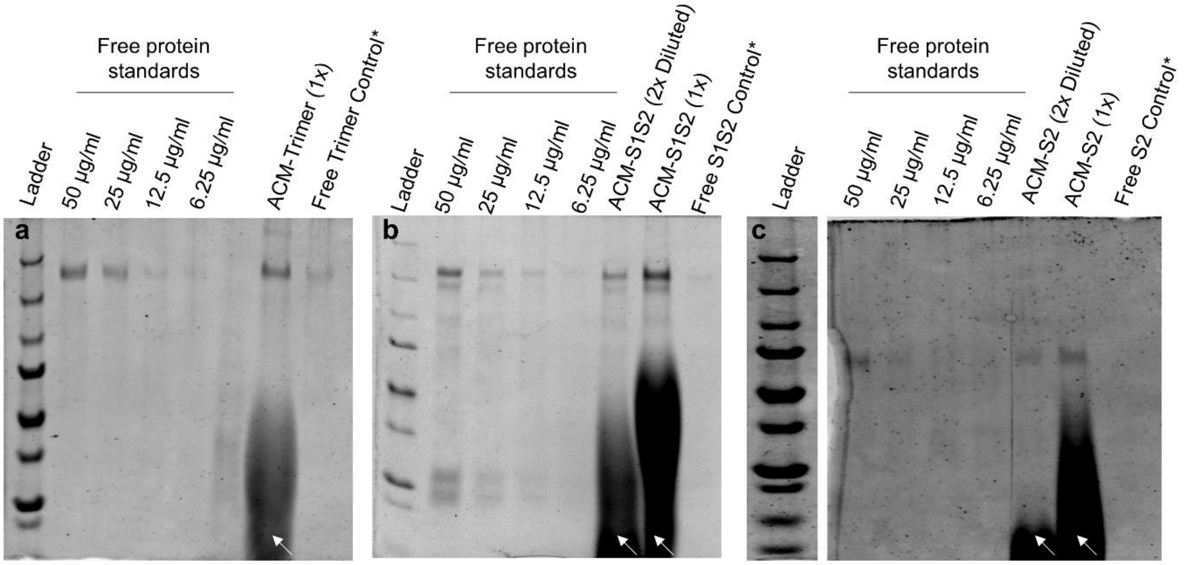

| Sample | Conc. (ug/ml) |
|---|---|
| ACM Trimer (1x) | 48.1 |
| Free Trimer Control | 19.8 |

| Sample | Conc. (ug/ml) |
|---|---|
| ACM-S1S2 (2x Diluted) | 43.7 |
| ACM-S1S2 (1x) | 49.2 |
| Free S1S2 Control | 7.5 |

| Sample | Conc. (ug/ml) |
|---|---|
| ACM-S2 (2x Diluted) | 25.5 |
| ACM-S2 (1x) | 25.8 |
| Free S2 Control | -0.7 |

**Supplementary Figure 3 | Assessing the amount of encapsulated protein by SDS-PAGE followed by SYPRO Ruby staining. a.** Trimer. **b.** S1S2. **c.** S2. *A parallel control experiment to estimate the amount of residual, non-encapsulated protein. White arrow: smear produced by ACM polymers.

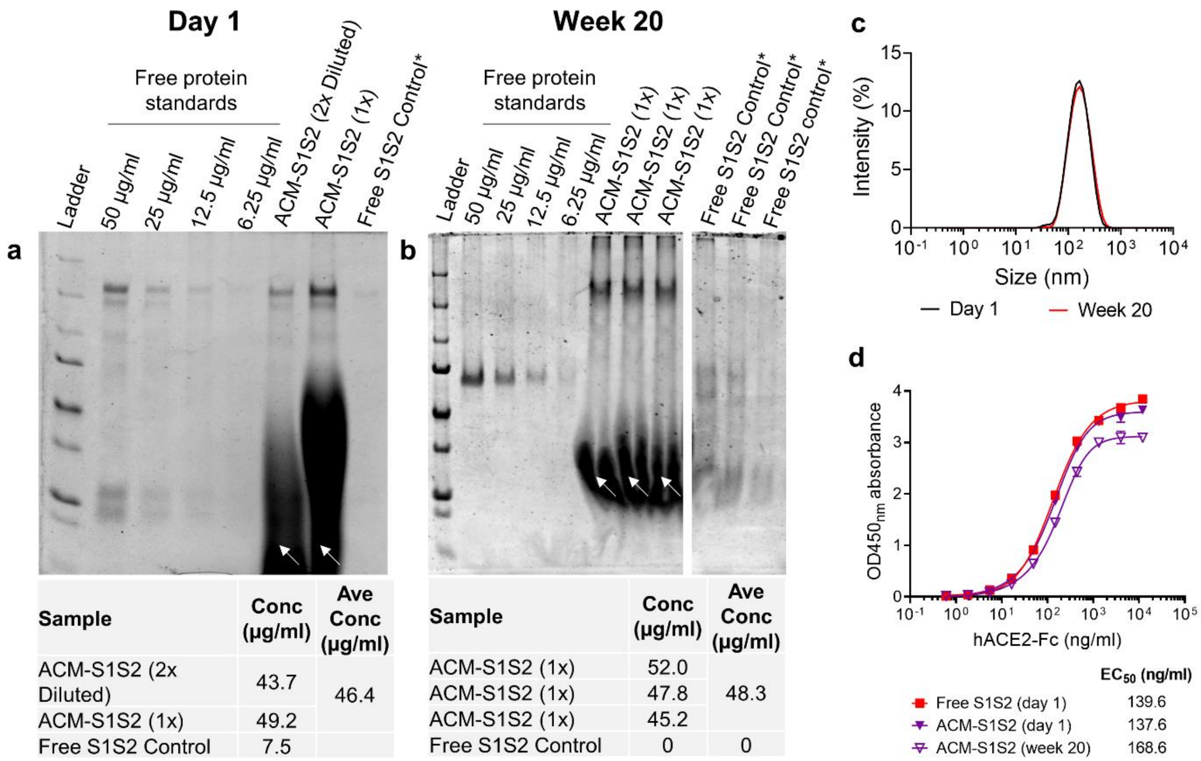

| Sample | Conc (μg/ml) | Ave Conc (μg/ml) |
|---|---|---|
| ACM-S1S2 (2x Diluted) | 43.7 | 46.4 |
| ACM-S1S2 (1x) | 49.2 | |
| Free S1S2 Control | 7.5 | |

| Sample | Conc (μg/ml) | Ave Conc (μg/ml) |
|---|---|---|
| ACM-S1S2 (1x) | 52.0 | 48.3 |
| ACM-S1S2 (1x) | 47.8 | |
| ACM-S1S2 (1x) | 45.2 | |
| Free S1S2 Control | 0 | 0 |

**Supplementary Figure 4 | Stability study of ACM-S1S2 at 4°C. a, b.** Quantity of ACM-encapsulated S1S2 on Day 1 and Week 20. ACM vesicles were lysed and protein was analyzed by SDS-PAGE and SYPRO staining. Day 1 concentration was calculated using free S1S2 protein standards; Week 20 concentration was calculated using free BSA standards due to lack of S1S2 protein. *A parallel control experiment to estimate the amount of residual, non-encapsulated protein. White arrow: smear produced by ACM polymers. **c.** DLS measurements of ACM polymersomes on Day 1 and Week 20 suggested no change in size and PDI of the ACM-S1S2 vesicles. **d.** ACE2 binding assay of ACM-S1S2 on Day 1 and Week 20 showed minimal loss of activity. Encapsulated S1S2 protein was released by lysing vesicles with Triton-X100.

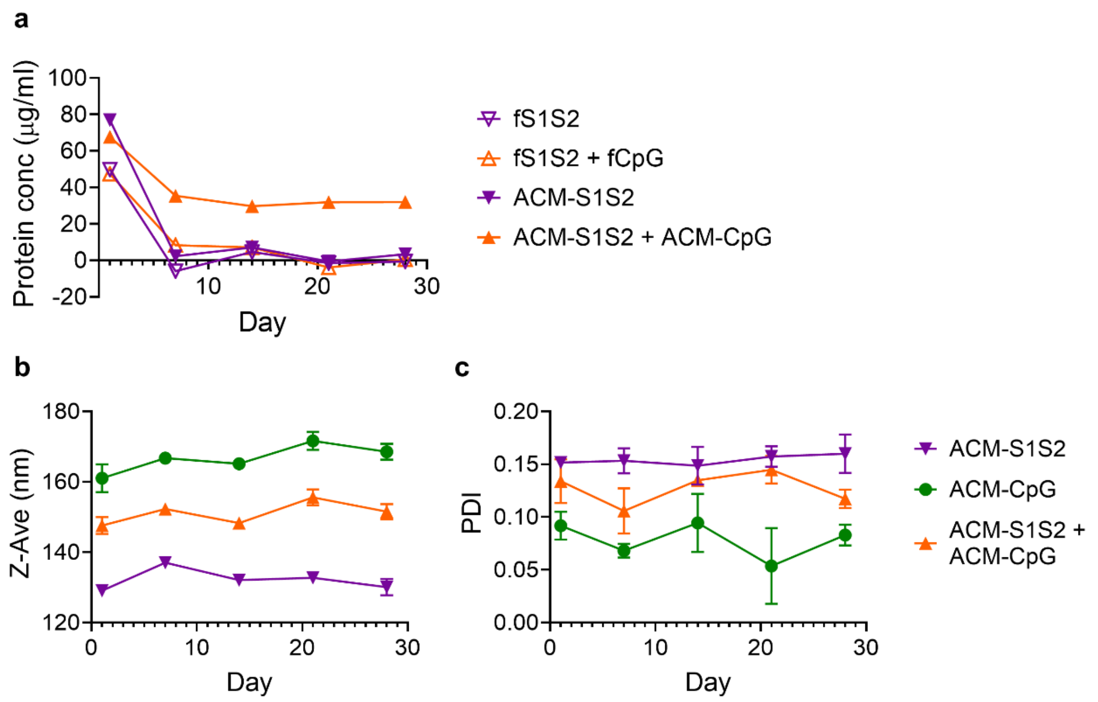

**Supplementary Figure 5 | Stability study of free S1S2, ACM-S1S2, free S1S2 + free CpG, and ACM-S1S2 + ACM-CpG at 37 °C for 28 days. a.** Amount of S1S2 protein present in different formulations over 28-day time course. **b, c.** Size and polydispersity (PDI) of ACM vesicles.

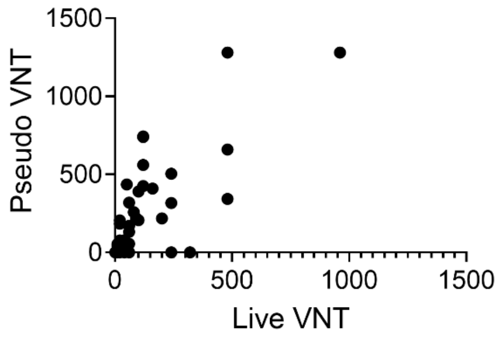

| | |
|---|---|
| Pearson r | |
| r | 0.8326 |
| 95% confidence interval | 0.7467 to 0.8911 |
| R squared | 0.6932 |
| P value | |
| P (two-tailed) | <0.0001 |
| P value summary | **** |
| Significant? (alpha = 0.05) | Yes |
| Number of XY Pairs | 75 |

**Supplementary Figure 6 | Correlation between pseudovirus and live virus neutralization tests.** Two-tailed Pearson correlation was performed between 75 pairs of data points from non-vaccinated and vaccinated mice from Day 54.

**a**

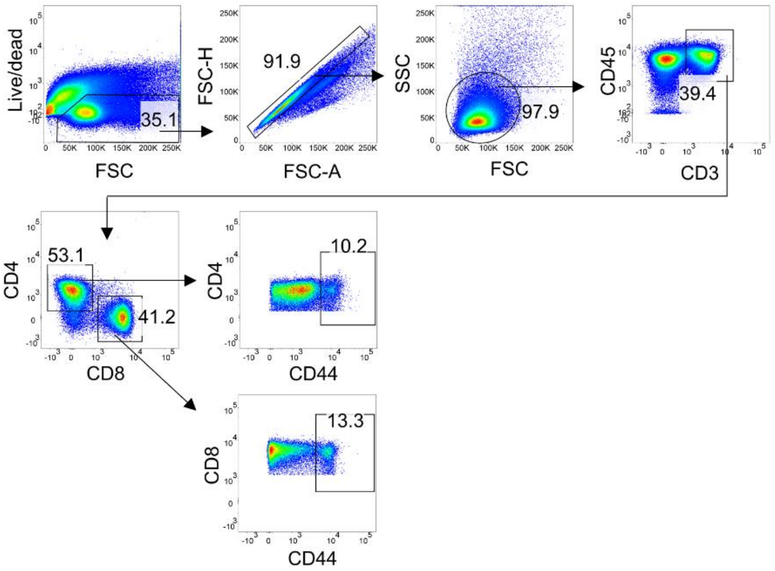
Live/dead
FSC
35.1
FSC-H
FSC-A
91.9
SSC
FSC
97.9
CD45
CD3
39.4
CD4
CD8
53.1
41.2
CD4
CD44
10.2
CD8
CD44
13.3

**b**

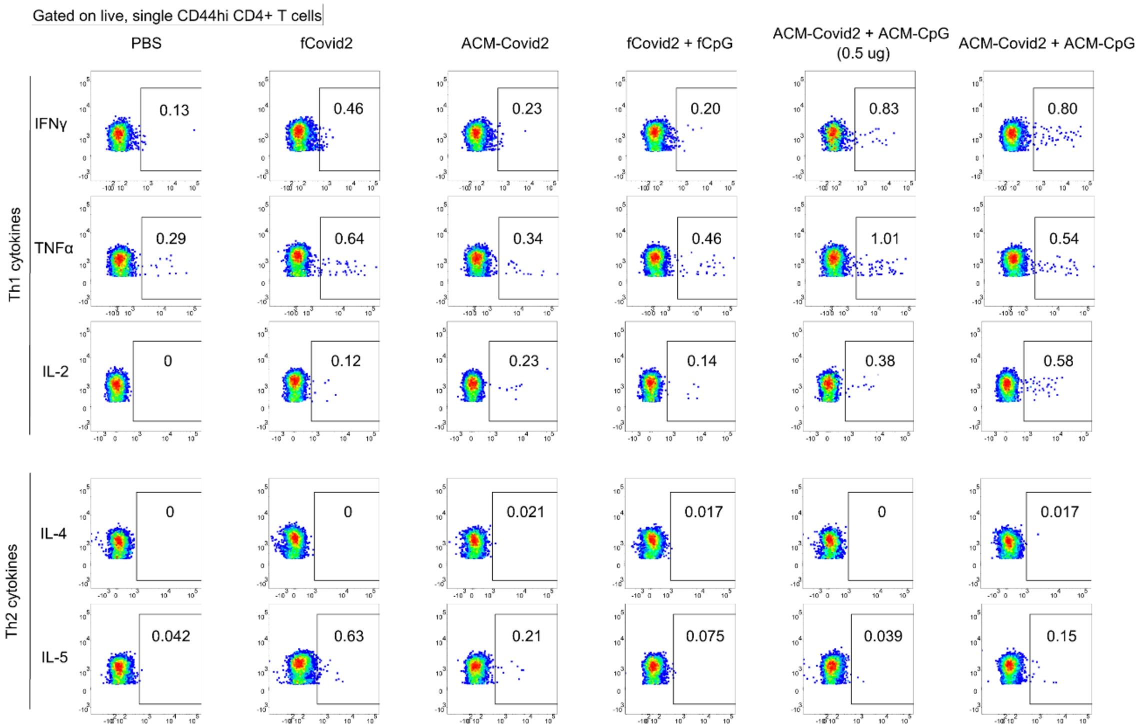
Gated on live, single CD44hi CD4+ T cells
PBS
fCovid2
ACM-Covid2
fCovid2 + fCpG
ACM-Covid2 + ACM-CpG (0.5 ug)
ACM-Covid2 + ACM-CpG
Th1 cytokines
IFNγ
0.13
0.46
0.23
0.20
0.83
0.80
TNFα
0.29
0.64
0.34
0.46
1.01
0.54
IL-2
0
0.12
0.23
0.14
0.38
0.58
Th2 cytokines
IL-4
0
0
0.021
0.017
0
0.017
IL-5
0.042
0.63
0.21
0.075
0.039
0.15

**c**

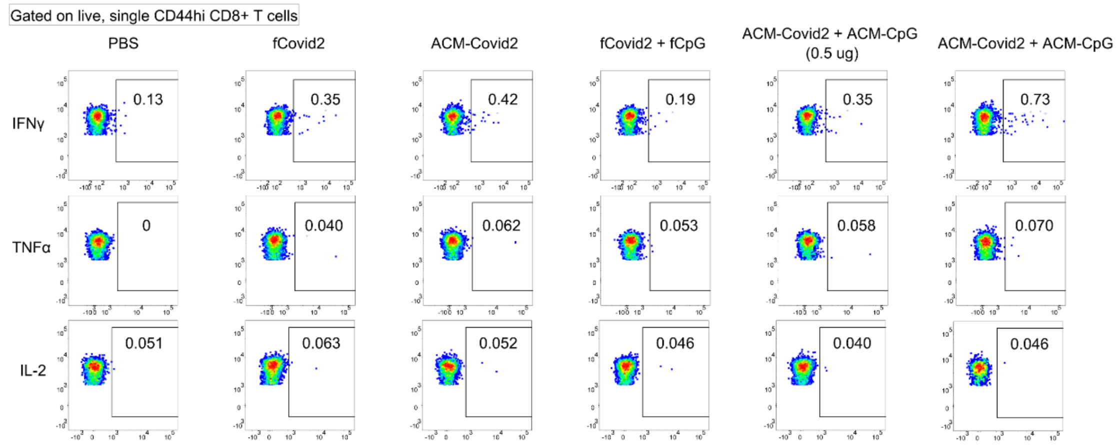
Gated on live, single CD44hi CD8+ T cells
PBS
fCovid2
ACM-Covid2
fCovid2 + fCpG
ACM-Covid2 + ACM-CpG (0.5 ug)
ACM-Covid2 + ACM-CpG
IFNγ
0.13
0.35
0.42
0.19
0.35
0.73
TNFα
0
0.040
0.062
0.053
0.058
0.070
IL-2
0.051
0.063
0.052
0.046
0.040
0.046

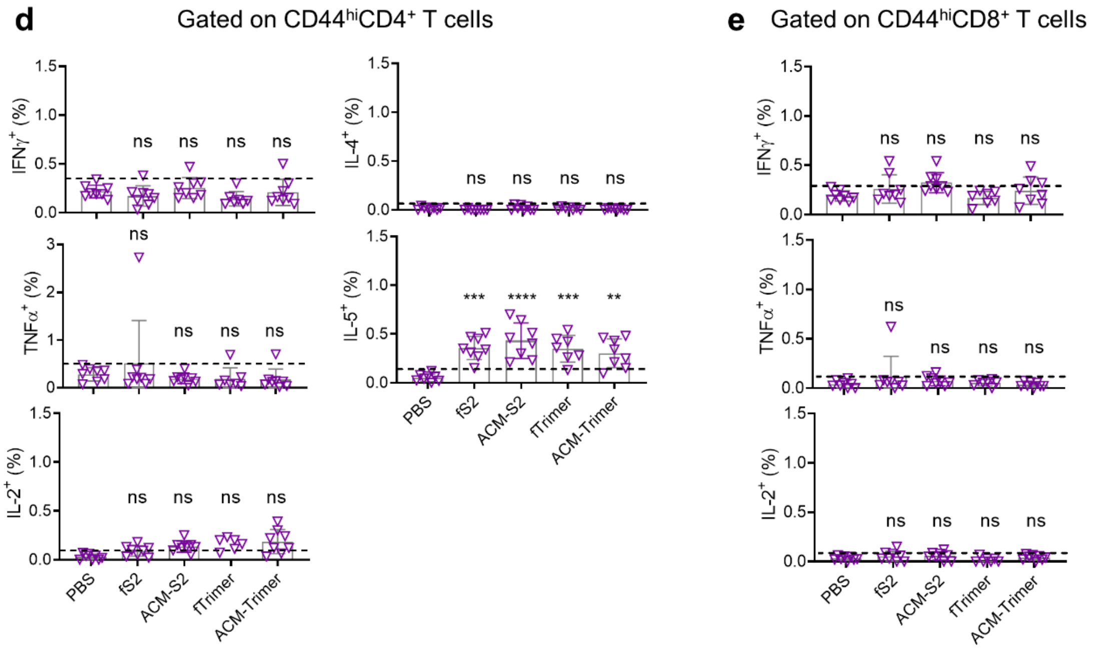

**Supplementary Figure 7 | Cytokine profiles of memory $CD4^+$ and $CD8^+$ T cells from immunized mice. a.** Gating strategy. $CD4^+$ and $CD8^+$ T cells possessing memory phenotype were identified by gating on the respective $CD44^{hi}$ subpopulations. **b, c.** Representative FCM plots from mice immunized with various S1S2 formulations, showing expression of IFNγ, TNFα, IL-2, IL-4 or IL-5 by $CD44^{hi}CD4^+$ or $CD44^{hi}CD8^+$ T cells in response to spike peptide stimulation. The number in each gate depicts percentage of cytokine-positive events. **d, e.** $CD4^+$ and $CD8^+$ T cell cytokine profiles of mice immunized with S2 or trimer formulations. Baselines (horizontal dashed lines) are assigned according to PBS controls and readings above them are considered antigen-specific. Statistical comparisons are made with respect to the PBS control using ordinary one-way ANOVA with Dunnett's multiple comparison. **: $P \leq 0.01$; ***: $P \leq 0.001$; ****: $P \leq 0.0001$; ns: not significant.
